# Myeloid cell-mediated killing of B-ALL by CD38 and CD20 IgA antibody variants is enhanced by CD47/SIRPα interference

**DOI:** 10.1101/2025.08.13.669665

**Authors:** M. Kowol, M. Lustig, A.M. Hartmann, I.C. van der Peet, M. Jansen, D. Winterberg, S. Krohn, N. Baumann, T. Rösner, S. Bendig, A.A. Wurm, M. Brüggemann, R. Burger, K. Schmitt, O. Valerius, F. Stölzel, L. Bastian, C.D. Baldus, M. Peipp, J.H.W. Leusen, D.M. Schewe, T. Valerius

## Abstract

Enhancing myeloid effector cell recruitment may improve immunotherapy by monoclonal antibodies – including that against acute lymphoblastic leukemia (ALL). To assess expression of target antigens in B-ALL, we compared mRNA profiling of 559 patient leukemia samples across 18 molecular subtypes with that of representative cell lines. The latter served as target cells to compare human IgG1 or IgA2 variants against CD19, CD20 or CD38 in antibody-dependent cellular phagocytosis (ADCP) by macrophages and antibody-dependent cell-mediated cytotoxicity (ADCC) by polymorphonuclear leukocytes (PMN). Interestingly, antibodies against broadly expressed CD19 were negligibly effective in mediating ADCP or ADCC. Antibodies against CD20 or CD38, the former variably expressed across subtypes, triggered ADCP by macrophages both as IgG1 and IgA2. However, PMN mediated ADCC against CD20 or CD38 was only observed with IgA2 variants, but not with respective IgG1 antibodies. Blocking the myeloid checkpoint molecule CD47 with a CD47 antibody or a soluble SIRPα-Fc fusion protein enhanced ADCP and ADCC by IgA2 antibodies. The binding site for SIRPα on CD47 contains an N-terminal pyroglutamate (pGlu), whose formation is catalyzed by glutaminyl-peptide cyclotransferase like (QPCTL). The direct involvement of pGlu in CD47/SIRPα interactions was shown by using engineered CD47 variants. Both CD47 and QPCTL were broadly expressed across BCP-ALL subtypes, indicating QPCTL inhibitors as additional therapeutic option. Importantly, the combination of anti-CD38 IgA2 and CD47 blockade was effective against xenografted B-ALL cells in human FcαRI (CD89) transgenic (tg) NXG mice. Together, these studies support the combination of anti-CD38 IgA2 with CD47 interference to improve myeloid effector cell recruitment for immunotherapy of B-ALL.

**Data sharing statement:** RNA-Seq data from BCP-ALL patients and B cells from healthy donors are available in the European Genome-Phenome Archive (EGA) accession numbers EGAS00001006107 and EGAS00001007305, respectively. Mass spectrometry data of analyzed proteins will be made available after manuscript acceptance on PRIDE - PRoteomics IDEntifications Database.

**Graphical abstract:** 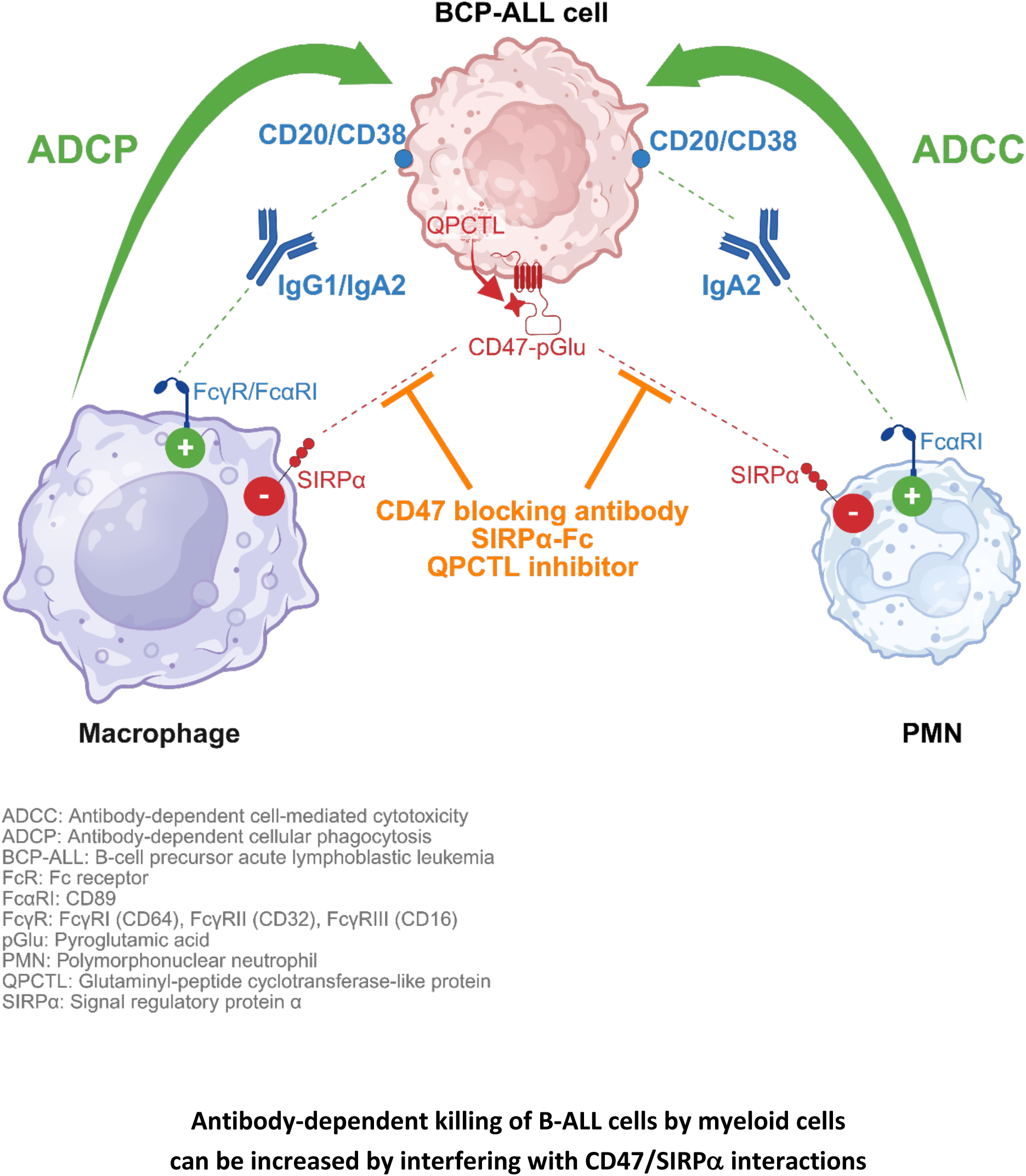

**Key points:**

- IgG1 and IgA2 antibodies against CD38 or CD20 were effective in recruiting macrophages for ADCP, but only IgA2 triggered ADCC by PMN
- Myeloid effector cell activation was enhanced by interfering with the CD47/SIRPα axis, especially when IgA2 antibodies were applied

## Introduction

Intensive chemotherapy has improved the outcome for patients with B-ALL, but often leads to treatment-related toxicities.^1^ Immunotherapy aims to reduce these toxicities while further improving outcomes.^2,3^ In addition to CAR T cells,^4^ antibody-based therapeutics - such as blinatumomab,^5^ rituximab (RTX)^6^ and inotuzumab-ozogamicin^7^ (targeting CD19, CD20 and CD22, respectively) – have become standard of care in adult ALL patients. We investigated approaches to improve myeloid effector cell recruitment in antibody-based therapy against different molecular subtypes of B-ALL.

Myeloid cells – such as monocytes/macrophages, granulocytes (polymorphonuclear cells, PMN) and myeloid-derived suppressor cells (MDSC) – are commonly found in the microenvironment of many solid tumors,^8,9^ and have been observed in close contact with B-ALL cells in the bone marrow.^10^ Across different tumor types, myeloid cell infiltrates are often associated with poor patients’ prognosis,^8,9^ but macrophages were also identified as important effector cell population for many monoclonal antibodies.^11,12^ For example, the efficacy of CD20 antibodies in syngeneic mouse models is mediated by macrophages,^13,14^ e.g. residing in the liver.^15^ Furthermore, studies in immunodeficient mice, which have functional myeloid but no lymphoid or NK cells, demonstrated therapeutic efficacy of antibodies against CD19 and CD20 against B cell malignancies.^16,17,18,19^ PMN are the most abundant myeloid cells, but are poorly recruited for tumor cell killing by human IgG1 antibodies.^20,21^ However, tumor cell killing by PMN is significantly enhanced when IgA2 antibodies engaging CD89 are employed instead of IgG1 antibodies,^22,23,24^ and when myeloid immune checkpoints are blocked.^25,26^

Similar to T cells, immune checkpoint molecules have also been described for myeloid cells, with CD47 on tumor cells and SIRPα on effector cells being the most established example.^27,28^ Checkpoint blockade by CD47 or SIRPα targeting molecules has been demonstrated to enhance the efficacy of therapeutic antibodies in many preclinical models,^28^ including T-and B-ALL.^29,30^ However, despite impressive early clinical results with RTX and the CD47 antibody magrolimab in B-NHL,^31^ the clinical development of magrolimab has recently been discontinued based on negative data in myeloid malignancies.^32^ Inhibition of QPCTL - the enzyme that catalyzes functionally relevant pGlu formation on many peptides and proteins including CD47 - was shown to modify the myeloid tumor cell infiltrate^33^ and to improve ADCP and ADCC against tumor cells.^34,35^

Here, we investigated the efficacy of CD47/SIRPα interference in combination with either human IgG1 or IgA2 antibodies against CD19, CD20, or CD38 in macrophage-mediated ADCP or PMN-mediated ADCC against B-ALL. In particular, anti-CD38 IgA2 with CD47 blockade evolved as a promising novel approach, which also proved effective in a xenograft model of human B-ALL in CD89 tg mice.

## Materials and Methods

### RNA profiling of patient samples and cell lines

RNA profiling of samples from a previously described cohort of 559 adult and pediatric B-ALL patients was performed as described.^36^ The same protocols were followed to analyze B-ALL cell lines.

### Cell lines

Human B-ALL cell lines were purchased from the German Collection of Microorganisms and Cell Cultures (DSMZ). Generation of gene-modified REH cells (luciferase-transduced REH^LUC^) is described in Suppl. Materials and Methods.

### Flow cytometric analyses

Flow cytometric analyses of cell lines were performed on a Navios flow cytometer (Beckmann Coulter) using protocols as previously described.^37,38^ To quantify the antigen density on the cell surface, the specific antibody binding capacity (SABC) was determined using QIFIKIT® (Agilent Technologies) according to the manufacturer’s instructions. Mouse blood samples were measured using flow cytometry (FACS Canto II) and analyzed using FACSDiva software.

### Therapeutic antibodies and generation of IgA2 variants

Human IgG1 antibodies were cetuximab (CTX; anti-EGFR; Merck), daratumumab (DARA; anti-CD38; Janssen Biotech), the CD20 antibodies RTX and obinutuzumab (OBZ) from Roche. Human IgA2.0 variants (abbreviated as IgA2), based on the variable regions of these antibodies, were generated as described.^23,39,37^ The CD19 antibodies are based on the variable regions of tafasitamab (TAFA; MorphoSys) and include an IgG1 wildtype,^18^ and an IgA2 variant, which was newly produced (Suppl. Material and Methods). The variable regions of the CD47 antibody magrolimab (hu5F9-G4, Gilead Sciences) were produced as an Fc-silent IgG2σ variant (Suppl. Material and Methods).^29^ A soluble SIRPα-Fc fusion protein (SIPRα-Fc) with silenced Fc (PGLALA), which blocks both human and murine CD47, was produced as described.^40^

### Isolation of human effector cells

PMN and peripheral blood mononuclear cells (PBMC) were isolated from the peripheral blood of healthy donors as described.^41^ Non-polarized (M0) macrophages were generated from monocytes stimulated with macrophage colony-stimulating factor (M-CSF) (PeproTech) for 11 to 14 days.

### Antibody-dependent effector cell-mediated assays

Antibody-dependent cellular phagocytosis (ADCP) was measured by live-cell imaging using the IncuCyte® System (Sartorius).^42^ Briefly, pHrodo (ThermoFisher) labeled target cells were added to M0 macrophages at an effector-to-target cell (E:T) ratio of 1:1. Antibodies were added as indicated and phagocytosis was measured every 20 min for a total of 6 h. Phagocytosis rates were quantified as red object counts per image (ROI).

PMN-mediated ADCC was measured in 3 h chromium-51 [^51^Cr] (Revvity) release assays or imaged with a Thunder RT widefield microscope (GE Healthcare) at an E:T ratio of 80:1.^37^ For all functional assays, antibodies were applied at a final concentration of 10 μg/ml (unless otherwise indicated), the anti-CD47 IgG2σ was used at 20 μg/ml. CTX and its IgA2 variant were used as isotype controls.

### In vivo studies

CD89 tg SCID and NXG mice^43^ and their non-transgenic (wt) littermates were bred and maintained at Janvier under a customized breeding agreement with the UMC. For the therapeutic study, REH^LUC^ cells were injected into NXG mice. Treatment with anti-CD38 IgA2 and soluble SIRPα-Fc fusion protein was given as explained in Supplemental files.

### Soluble CD47-Fc fusion proteins and QPCTL inhibitor

Soluble Fc fusion proteins with native human CD47 (CD47-wt) and a Q1A variant (CD47-Q1A) were generated and functionally tested as described (Suppl. Materials and Methods).

The small molecule QPCTL inhibitor SEN177 (Sigma-Aldrich) was dissolved in DMSO (Thermo Fisher) and used as described.^44^

### Statistical analyses

Flow cytometric analyses and functional in vitro assays were performed in at least three independent experiments with effector cells from different donors. Depicted values are means +/- standard error of the mean (SEM). Statistical significance was determined using one-or two-way ANOVA with Bonferroni correction. p-values < 0.05 were considered significant. Graphical and statistical analyses were performed using GraphPad Prism 10.

Statistical Analyses for RNA-Seq data were performed in the R-package “ggpubr”^45^, using one-way ANOVA for multiple comparisons of gene expression means and t-test for pairwise comparison of means (leukemic vs healthy B cell gene expression).

### Animal welfare

The animal studies underwent review and were granted approval by the Animal Ethical Committee of the UMC Utrecht (AVD number: 115002021 15442).

### Ethical approval

In accordance with the Declaration of Helsinki, the Ethics Committees of the respective institutions approved all studies with isolated effector cells from healthy human blood donors, who gave written informed consent before analyses (University Medical Center Utrecht: 07-125/O, Christian-Albrecht University Kiel: D 563/18).

Additional methodological details are provided in the Supplement.

## Results

### Correlation between mRNA-Seq data of B-ALL cell lines and patient samples from corresponding molecular subtypes

B-ALL has long been classified according to its immunophenotype^46^ - resembling physiological stages of B cell differentiation.^47,48^ More recently, mRNA profiling emerged as a diagnostic alternative to more precisely classify B-ALL samples based on their transcriptional activity.^49^ The ALLCatchR tool allocates 21 gene-expression defined molecular B-ALL subtypes from RNA-Seq profiles with high accuracy.^36^ To identify potential target antigens for immunotherapy, we tested mRNA expression levels of antigens against which approved antibody therapeutics are available across an adult cohort representing 18 of these subtypes (Figure 1A). In search for appropriate leukemia cell lines for functional studies, we generated mRNA profiles from eight candidate cell lines representing seven molecular entities, and analyzed if their gene expression profiles matched with respective patient samples. Uniform manifold approximation and projection (UMAP) clustering of all batch-corrected samples according to the LASSO-selected subtype-specific gene sets used in the ALLCatchR resulted in a clear separation of molecular subtypes (Figure 1B). Except for cell line TOM-1 (*BCR::ABL1*-positive), the other seven tested BCP-ALL cell lines clustered with the respective patient samples of their corresponding subtypes. These results support that the selected cell lines can serve as representative models.

**Figure 1.**
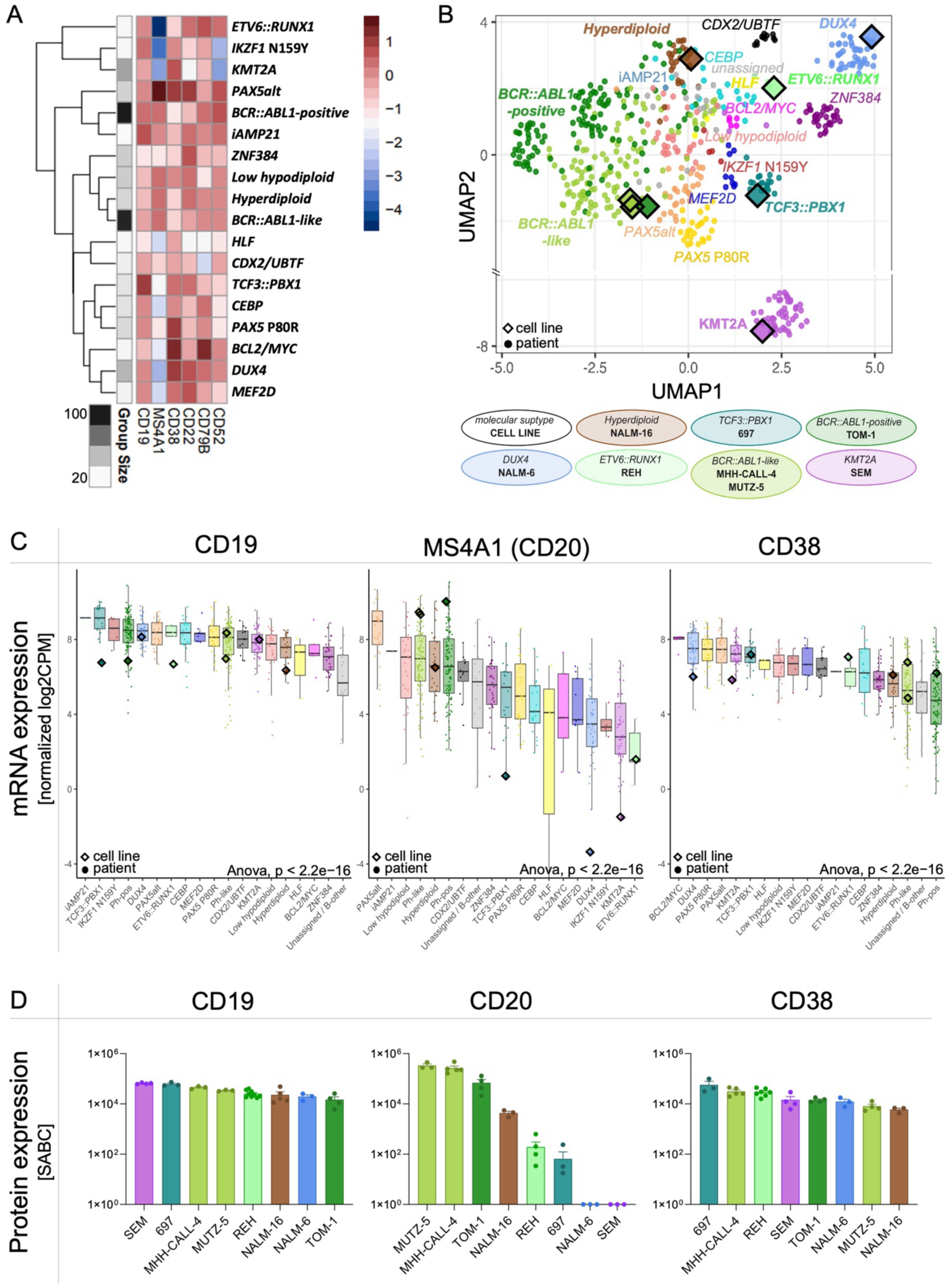
Correlation between mRNA-Seq data of B-ALL cell lines and patient samples from corresponding molecular subtypes. **(A)** Comparison of gene expression of genes of interest. Log2-fold changes of the mean expression per subtype vs mean expression of the cohort were scaled gene-wise and plotted using heatmap package.^53^ **(B)** Uniform manifold approximation and projection (UMAP) plot of molecular subtypes derived from mRNA-Seq data from a B-ALL patient cohort (n = 559 patient samples, as published^36^) compared to eight selected cell lines. One dot represents one patient, each rhomb one cell line. Individual colors represent different molecular subtypes. **(C)** CD19, MSA41 (CD20), and CD38 mRNA expression, depicted as normalized log2CPM, in patient leukemia samples (dots) and selected cell lines (rhombs) classified by molecular subtypes. The groups are ordered by descending median expression levels. The lower and upper hinges correspond to the first and third quartiles (the 25th and 75th percentiles). **(D)** CD19, CD20, and CD38 protein expression on the cell surface of B-ALL cell lines as analyzed by quantitative flow cytometry. Depicted are specific antibody binding capacities (SABC) as mean values ± SEM of at least three experiments (indicated as dots). The cell lines are ordered by their expression levels, the bar colors correspond to the respective molecular subtypes (shown in B and C).

Next, we assessed mRNA expression levels of three potential target antigens for myeloid effector cell recruitment (CD19, CD20 and CD38) across the respective subtypes in more detail (Figure 1C). We found CD19 broadly expressed both within patient samples and cell lines (Figure 1C, left). Expression levels of MS4A1 (CD20) were found to be more subtype-related, with particularly low levels in *ETV6::RUNX1* samples and high levels in PAX5alt samples (Figure 1C, center). Among the cell lines, MHH-CALL-4 and MUTZ-5 (both *BCR::ABL1*-like) and TOM-1 (*BCR::ABL1*-positive) showed the highest CD20 mRNA expression levels, while others like NALM-6 (*DUX4*), SEM (*KMT2A*) and 697 (*TCF3::PBX1*) contained very low levels. CD38 was broadly expressed by all subtypes with lowest levels in *BCR::ABL1*-like and *BCR::ABL1-*positive cell lines (Figure 1C, right). For all three antigens, significant variability between individual patient samples was observed within all subgroups (multi-comparison ANOVA, p < 2.2×10^-16^), which was particularly large for CD20. Since previous studies have shown that myeloid effector cells require rather high antigen expression levels to mediate tumor cell killing,^50,51,52^ we measured target antigen expression on the previously identified candidate B-ALL cell lines. Here, all B-ALL cell lines expressed similar levels of CD19 and CD38, while CD20 expression was more variable and absent on some cell lines (Figure 1D). Subsequently CD19, CD20 and CD38 served as target antigens for ADCP by macrophages and ADCC by PMN.

### Myeloid cell-mediated tumor cell killing with CD19, CD20, and CD38 antibodies of IgG1 and IgA2 isotypes

To investigate whether myeloid effector cell activation against B-ALL can be improved by IgA2 compared to IgG1 antibodies, IgA2 variants of clinically approved IgG1 antibodies against CD19, CD20 and CD38 were generated using the variable regions of TAFA, OBZ and DARA, respectively (Figure 2A). Antigen binding was similar for IgG1 and IgA2 variants (Figure 2B). Against CD20 also a matched set of RTX variants was analyzed (Suppl. Figure 1A, B). Based on the antigen expression levels observed in Figure 1D, we tested cell lines with high expression levels for each antigen in myeloid cell-mediated killing assays. Thus, SEM (*KMT2A*) and 697 (*TCF3::PBX1*) served as target cells for CD19, 697 and REH (*ETV6::RUNX1*) for CD38, and MUTZ-5 (*BCR::ABL1*-like) and MHH-CALL-4 (*BCR::ABL1*-like) for CD20. While both the IgG1 and IgA2 variants of the CD19 and CD38 antibodies were similarly effective in triggering ADCP by macrophages (Figure 2C), the glyco-engineered IgG1 antibody OBZ against CD20 performed better than its IgA2 variant (Figure 2C). On the other hand, only the IgA2 but not the IgG1 variants of CD20 and CD38 antibodies mediated ADCC with PMN as effector cells (Figure 2D). Compared to RTX IgG1, OBZ IgG1 showed enhanced ADCP of MHH-CALL-4 (but not MUTZ-5), while OBZ IgA2 was consistently superior to RTX IgA2 in PMN-mediated ADCC (Suppl. Figure 1C). Against CD19, neither IgG1 nor IgA2 antibodies recruited PMN for ADCC, while significant ADCP was observed against some, but not all cell lines (Figure 2C, D, Suppl. Figure 1D). Thus, anti-CD38 and anti-CD20 were effective as IgG1 and IgA2 in ADCP by macrophages, but only IgA2 variants against both antigens mediated ADCC by PMN. Since IgG1 dependent leukemia cell killing is well documented,^18,19,29,30,54^ focused on IgA2 in our further experiments.

**Figure 2.**
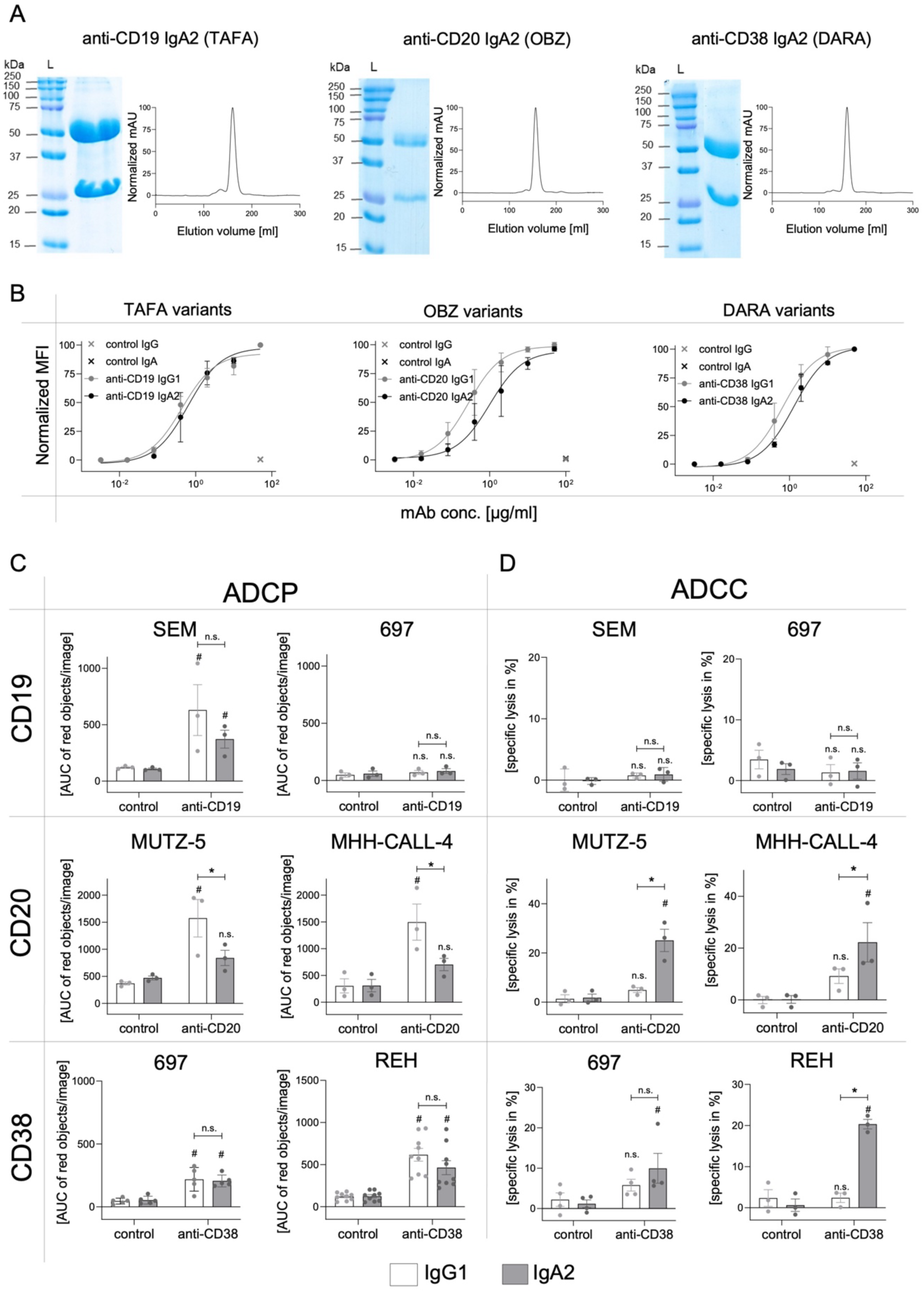
Myeloid cell-mediated tumor cell killing with CD19, CD20, and CD38 antibodies of IgG1 and IgA2 isotypes. **(A)** SDS-PAGE under reducing conditions and size exclusion chromatography of the IgA2 antibodies (TAFA, tafasitamab; OBZ, obinutuzumab; DARA, daratumumab) under native buffer conditions. Representative chromatography images show the isolated peak fractions normalized to maximum absorption. **(B)** Binding capacities of the different IgA2 antibodies (TAFA, OBZ, DARA) on 697 cells (CD19 and CD38) and on MUTZ-5 cells (CD20) at increasing concentrations up to 50 µg/ml. Respective antibodies were detected by FITC labelled goat anti-human kappa light chain F(ab’)_2_ fragments. Shown are the normalized MFI values +/- SEM of three independent experiments. **(C+D)** ADCP by M0 macrophages **(C)** and ADCC with PMN **(D)** using the indicated B-ALL cell lines as targets. Antibodies were used at a concentration of 10 µg/ml. ADCP results are shown as area under the curve (AUC) ± SEM of phagocytosed cells (red objects per image) over 6 h. ADCC results are shown as mean specific lysis values ± SEM. All results represent at least 3 independent experiments with effector cells from different donors, each performed in triplicates. Data were analyzed by two-way ANOVA, and significant differences (p ≤ 0.05) between IgG1 and IgA2 (*) or specific vs control antibodies (control) (#) are indicated. n.s., not significant.

### CD47 expression on B-ALL cells and its role as immune checkpoint in ADCC and ADCP in vitro

Myeloid cell mediated tumor cell killing is regulated by the immune checkpoint molecules CD47/SIRPα.^55,28^ Thus, we assessed CD47 mRNA expression across the molecular B-ALL subtypes and correlated its expression to our selected cell lines. As shown (Figure 3A), CD47 is broadly expressed at high mRNA levels in all B-ALL subtypes and in our cell lines, reflected by cell surface expression on cell lines (Figure 3B). An Fc-silent CD47 blocking antibody based on the variable regions of magrolimab (anti-CD47 IgG2σ^29^) was used to investigate the impact of CD47 blockade on ADCC and ADCP by IgA2 antibodies. Importantly, CD47 blockade alone (in the absence of a tumor targeting antibody) did not trigger ADCP or ADCC in vitro (Figure 3C, D) – supporting that activating signals on effector cells e.g. through ITAM-containing FcR are required.^55,56^ Importantly, both ADCP and ADCC were significantly enhanced in the presence of IgA2 variants of CD20 and CD38 antibodies (Figure 3C, D, Suppl. Figure 2A). In contrast, the CD47 blocking antibody combined with CD19 IgA2 did not enhance ADCC and only enhanced ADCP in two out of six tested cell lines (Figure 3C, D, Suppl. Figure 2B). Representative ADCP and ADCC experiments are visualized in Figure 3E and F.

**Figure 3.**
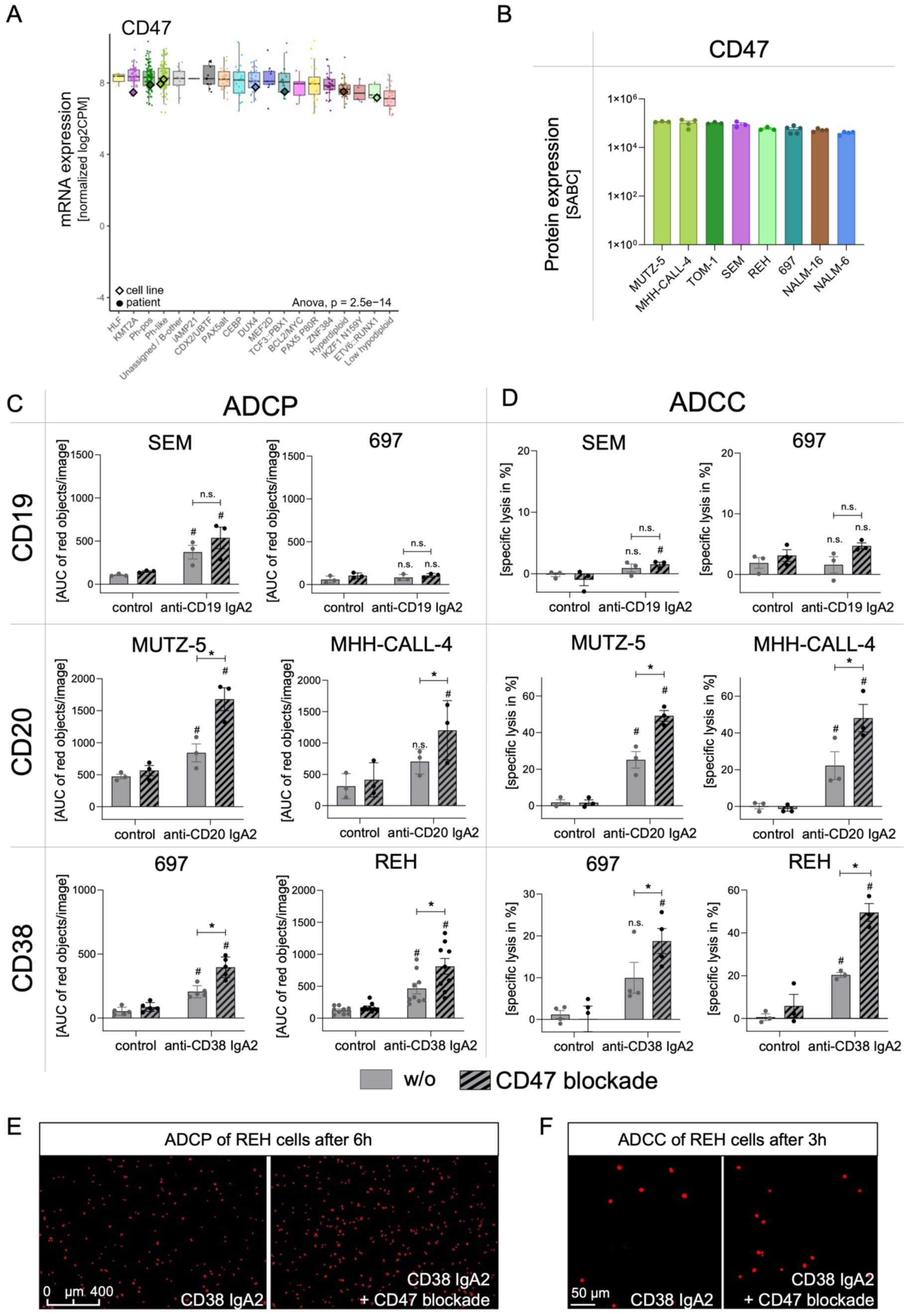
CD47 expression on B-ALL cells and enhanced efficacy of IgA2 mediated ADCP and ADCC in combination with CD47 blockade. **(A)** CD47 mRNA expression (normalized log2CPM) on patient leukemic cells (dots) and selected cell lines (rhombs), grouped by the molecular subtypes. **(B)** CD47 protein expression levels on the cell surface of B-ALL cell lines as analyzed by quantitative flow cytometry. Results are depicted as mean specific antibody binding capacity (SABC) ± SEM of 3 independent experiments. The bar colors correspond to the molecular subtypes as shown in A. **(C+D)** B-ALL cell lines served as target cells in ADCP **(C)** and ADCC **(D)** experiments as described in Materials & Methods. IgA2 variants of tafasitamab (CD19), obinutuzumab (CD20), and daratumumab (CD38) antibodies were used at 10 µg/ml in the presence or absence of an anti-CD47 IgG2σ antibody (20 µg/ml). The results are presented in the same manner as in Figure 2C+D. Data were analyzed by two-way ANOVA, and significant differences (p ≤ 0.05) between IgA2 alone or in combination with CD47 blockade (*) or specific vs control antibodies (control) (#) are indicated. n.s., not significant. **(E)** Microscopic images of ADCP depicting phagocytosed REH cells at 6 h in the presence of anti-CD38 IgA2 alone (10 µg/ml) and in combination with anti-CD47 IgG2σ (20 µg/ml). **(F)** Visualization of a representative ADCC experiment using live-cell imaging, showing dead REH cells after 3 h. Antibody concentrations were as in (D).

### CD47 blockade improves therapeutic efficacy of an CD38 IgA2 antibody in B-ALL in vivo

CD47 blockade has been shown to enhance the in vivo activity of IgG1 antibodies targeting several antigens,^55,28^ including CD19 in B-ALL^30^ and CD38 in T-ALL^29^. However, the combination of CD47/SIRPα blockade and IgA antibodies has been investigated in more limited settings,^25,39,37,26^ so far not including ALL. Since CD38 antibodies are increasingly investigated in ALL patients^57^, we investigated whether an IgA2 variant of DARA would be effective against B-ALL in CD89 tg NXG mice. In a pilot experiment, we compared the engraftment of REH^LUC^ cells in CD89 tg SCID and NXG mice (Suppl. Figure 3A, B). Since outgrowth was less heterogeneous in NXG than in SCID mice and the former are more commonly used in ALL research, 8-week-old female NXG mice were used for the therapeutic study.

Six groups of NXG mice (CD89 tg or WT as indicated) received 4 x 10^6^ REH^LUC^ cell i.v. and were treated i.p. with vehicle, SIRPα-Fc, anti-CD38 IgA2, or the combination of both as indicated in Figure 4A. All mice received pegylated (peg-) G-CSF s.c. on days −2 and +5, which is known to enhance tumor cell killing by CD89^58^. Based on previous experience with IgA2 antibodies in vivo^22,23^, mice were treated with anti-CD38 IgA2 at the indicated time points (Figure 4A). To block CD47/SIRPα interactions in vivo, we used an engineered human SIRPα-Fc with high affinity for both mouse and human CD47 fused to a silent Fc^26^, that increased ADCC and ADCP similar to the previously employed CD47 blocking antibody (Suppl. Figure 4A, B). The SIRPα-Fc was injected i.p. at days 0 and +12 to test for its impact on leukemia cell outgrowth alone or in combination with anti-CD38 IgA2. Ex vivo analyses on blood samples showed that monocyte and granulocyte counts were similar in all treatment groups (Suppl. Figure 5A). As previously described,^43^ CD89 expression was found on neutrophils in CD89 tg but not in wt mice (Suppl. Figure 5B). Human IgA on granulocytes was not detected at any time point (Suppl. Figure 5C), while SIRPα-Fc on granulocytes was observed until day +58 (Suppl. Figure 5D).^59^ With respect to therapeutic efficacy, BLI images showed that mice in the control groups 1 (vehicle) and 2 (anti-CD38 IgA2 in non-tg mice) were the first to develop tumors. Importantly, anti-CD38 IgA2 alone was effective in CD89 tg but not in wt mice (Figure 4B, C and Suppl. Figure 5E, upper panels). As expected from previous results,^29,30^ SIRPα-Fc alone was effective in tg and wt mice, and outperformed vehicle control and IgA2 monotherapy (Figure 4B, C, D). The combination treatment of SIRPα-Fc and anti-CD38 IgA2 in tg mice resulted in the highest number of tumor free mice (Figure 4B) and the greatest prolongation of survival (Figure 4C), although the survival difference was not statistically significant compared to treatment with SIRPα-Fc alone (Figure 4C). Nevertheless, a significantly higher percentage of mice were tumor free as assessed by BLI after combination therapy in the tg compared to the wt combination group and compared to the SIRPα-Fc -alone group. BLI data showed that the combination of anti-CD38 IgA2 with SIRPα-Fc was more effective in CD89 tg mice compared to wt mice (Suppl. Figure 5E, lower panels). Together, these in vivo results support the therapeutic efficacy of CD47 blockade in combination with anti-CD38 IgA2 in the treatment of B-ALL.

**Figure 4.**
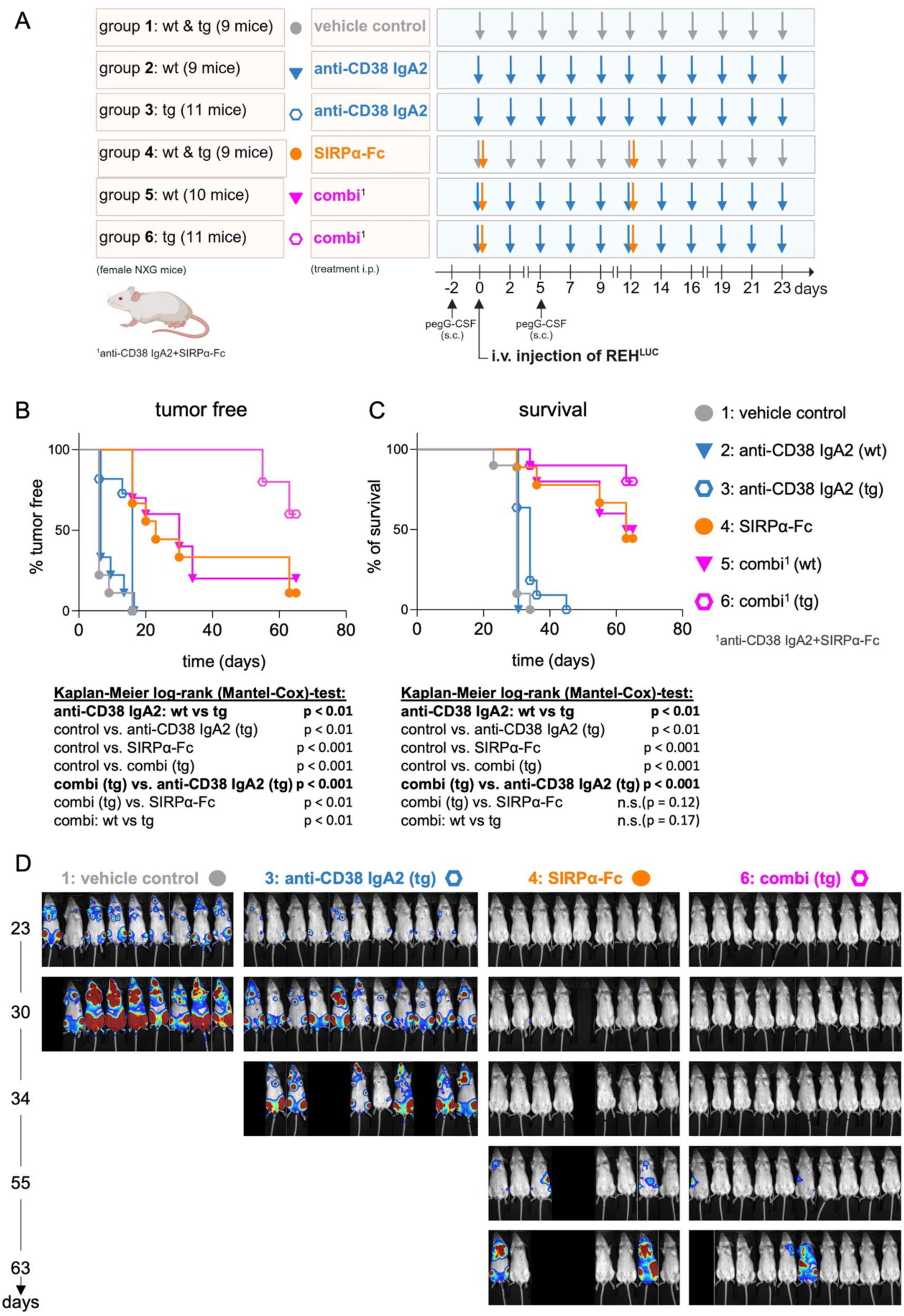
CD47 blockade improves the therapeutic efficacy of anti-CD38 IgA2 in a xenograft B-ALL leukemia model in FcαRI (CD89) transgenic NXG mice. **(A)** Treatment schedules of CD89 transgenic (tg) and wild type (wt) NXG mice (9-11 mice per group). Mice were injected with 4×10^6^ REH^LUC^ cells and treated with anti-CD38 IgA2 (blue arrows) and Fc silent SIRPα-Fc fusion protein (orange arrows) as indicated. Mice were subjected to bioluminescence (BLI) imaging 1-2 per week. Once a week, blood was collected for flow cytometric analyses. **(B)** Percentage of tumor free mice as assessed by BLI over 65 days presented as Kaplan-Meier curves. Data were analyzed by log-rank (Mantel-Cox) test. n.s., not significant. **(C)** Kaplan-Meier survival curves of mice as assessed over 65 days. Data were analyzed by log-rank (Mantel-Cox) test and p-values are shown. n.s., not significant. **(D)** Outgrowth of REH^LUC^ cells in mice under different therapeutic conditions as presented by front scan BLI images at the indicated days.

### QPCTL-mediated posttranslational pyroglutamate formation of CD47 impacts the efficacy of CD47/SIRPα interactions and can be targeted by small molecule inhibitors

The functional activity of CD47 is regulated by its enzymatic modifier QPTCL.^34,35^ mRNA profiling revealed that QPTCL expression is more strongly upregulated than that of CD47 in B-ALL compared to normal B cells (Figure 5A). QPCTL mRNA expression could also be confirmed across all molecular B-ALL subtypes and in our respective cell lines (Figure 5B). The Golgi-resident enzyme QPCTL catalyzes N-terminal pGlu formation,^60,34,35^ which is part of the CD47 binding pocket for SIRPα, as known from crystallographic analyses (Figure 5C).^61^ However, on a molecular level the direct involvement of pGlu in the CD47/SIRPα interaction has not been confirmed yet. We generated human CD47 Fc-silent fusion proteins (IgG1σ-Fc),^62^ that contain either a native CD47 ectodomain with N-terminal pGlu (CD47-wt) or a mutated version with alanine at the N-terminus (CD47-Q1A) (Figure 5D). Amino acid exchange and the posttranslational modification of glutamate to pGlu were confirmed by mass spectrometry. Here, the two different N-terminal fragments could be identified as separate peaks (Figure 5E). Importantly, the two CD47-Fc fusion proteins displayed clear differential binding to recombinant SIRPα in ELISA experiments with significantly reduced binding of the mutated CD47-Q1A molecule compared to CD47-wt (Figure 5F). Consistent with the expression of SIRPα on macrophages (Suppl. Figure 6A), binding of CD47-Q1A to macrophages was significantly reduced compared to CD47-wt binding (Figure 5G), supporting that pGlu is essential for CD47/SIRPα interactions. Incorrect folding of the mutated CD47 ectodomain is unlikely since binding of the pGlu-independent CD47 antibody B6H12 (binding total CD47) was similar for CD47-wt and CD47-Q1A (Suppl. Figure 6B, left). In contrast, the pGlu-dependent CD47 antibody CC2C6^34^ hardly interacted with the mutated CD47-Q1A, confirming that binding of this antibody depends on the presence of pGlu (Suppl. Figure 6B, right).

**Figure 5.**
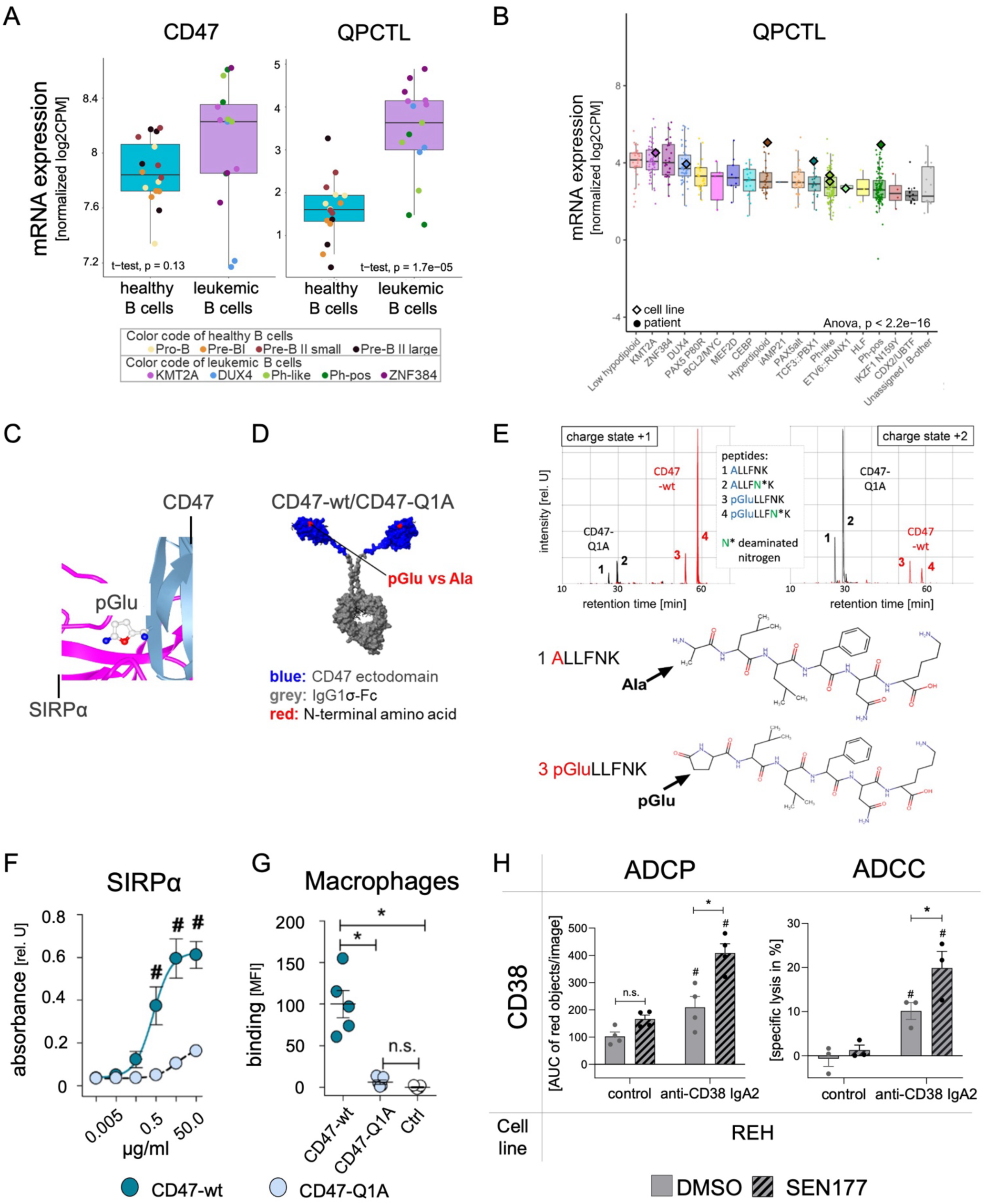
QPCTL-mediated pyroglutamate (pGlu) formation on the CD47 ectodomain mediates SIRPα binding and inhibits antibody-dependent tumor cell killing by myeloid cells. **(A)** CD47 (left panel) and QPCTL (right panel) mRNA levels of 15 samples of leukemic cells (violet bars) in comparison to 16 samples of healthy B cells of early developmental B cell stages (blue bars). Dots represent individual samples of a previously described cohort^36^ as indicated by their color codes. Data were analyzed by student t-test. **(B)** QPTCL mRNA expression (normalized log2CPM) on patient leukemic cells (dots) and selected cell lines (rhombs) grouped by the different molecular subtypes. The lower and upper hinges correspond to the first and third quartiles (the 25th and 75th percentiles). **(C)** Cartoon from the crystal structure (2JJT^61^) of the CD47 ectodomain (blue) containing pGlu within the binding pocket in complex with the SIRPα ectodomain (purple). **(D)** Structures of CD47-Fc fusion proteins generated by homology modeling. CD47-wt contains a native CD47 ectodomain with N-terminal pGlu, which is mutated to an N-terminal alanine (Ala) in CD47-Q1A. Both proteins are linked to an IgG1-Fc lacking Fcγ receptor binding (IgG1σ-Fc). **(E)** After gel electrophoresis, relevant protein bands (approx. 42 kDa) were analyzed by mass spectrometry to identify N-terminal pGlu in CD47-wt or alanine in CD47-Q1A. N-terminal CD47 fragments ALLFNK and ALLFN*K (* for deaminated nitrogen) of CD47-Q1A (peaks 1 and 2) and pGluLLFNK and pGluLLFN*K of CD47-wt (peaks 3 and 4) were identified at charge state +1 (left) and charge state +2 (right). Chromatograms were illustrated with RawMeat (Vast Scientific, Inc.). The peptide structures are shown below. **(F)** Differential binding of CD47-wt and CD47-Q1A to recombinant SIRPα. SIRPα (0.5 µg/ml) was coated to ELISA plates, CD47 fusion proteins were added at indicated concentrations, and binding detected by mouse HRP-conjugated anti-human IgG (Fc-specific). Mean absorbance values ± SEM of three independent experiments are shown. # indicates statistical differences between CD47-wt and CD47-Q1A at the indicated concentrations (p < 0.05 by two-way ANOVA with Bonferroni correction). **(G)** Differential binding of CD47 fusion proteins (50 µg/ml) to M0 macrophages as analyzed by indirect immunofluorescence using FITC-conjugated anti-human IgG F(ab‘)_2_ fragments for detection. Mean fluorescence intensity (MFI) values of five independent experiments are shown. * indicates statistically significant differences between CD47-wt and CD47-Q1A or the control (Ctrl: detection antibody alone) (p < 0.05 by one-way ANOVA with Bonferroni correction). n. s., not significant. **(H)** REH cells were treated with QPCTL inhibitor SEN177 in DMSO or DMSO alone (both 10 μM, 72 h). The IgA2 variant of daratumumab (10 µg/ml) was used as targeting antibody in ADCC (upper panel) and ADCP (lower panel). For ADCC, PMN at an E:T cell ratio of 80:1 were applied. ADCC results are shown as mean lysis rates in percent ± SEM of at least 3 independent experiments with effector cells from different donors. For ADCP, results are shown as area under the curve (AUC) ± SEM of phagocytosed REH cells (red objects per image, ROI) over 6 h. ADCP data represent 4 independent experiments using cells from different donors, each performed in triplicates. Data were analyzed by two-way ANOVA, and significant differences (p ≤ 0.05) between DMSO and SEN177 treated cells (*) or between the specific and control antibody (#) are indicated.

The QPTCL catalyzed posttranslational pGlu formation of CD47 can be inhibited by small molecules.^63^ To investigate the functional impact of QPCTL in the context of CD38-targeted therapy for B-ALL, REH cells were treated with the QPCTL inhibitor SEN177. As shown in Suppl. Figure 6C, soluble SIRPα binding was reduced on SEN177 treated REH cells. Consequently, QPCTL inhibition by SEN177 improved both ADCC and ADCP of REH cells by anti-CD38 IgA2 compared to the respective control cells (Figure 5H). Overall, these data support the involvement of QPCTL in regulating CD47/SIRPα interactions in B-ALL cells and suggest small molecule QPCTL inhibitors as another therapeutic approach to interfere with this myeloid checkpoint.

## Discussion

ALL has emerged as a model disease for novel immunotherapeutic approaches, with the first bispecific T cell engager (blinatumomab^64^), the first CAR T cell product (tisagenlecleucel) and the second ADC (inotuzumab-ozogamicin^7^) being approved for B-ALL.^2,1^ Nevertheless, the development of conventional monoclonal antibodies is lacking behind in ALL, although many B cell restricted antigens have been identified. Thus, we analysed mRNA expression levels across 18 previously defined molecular B-ALL subgroups^36^ for potential target antigens, against which antibody-based products are clinically approved.^65,54^ We then focused on CD19, CD20 and CD38 and investigated how myeloid effector cell recruitment by antibodies against these target antigens could be improved.

Data presented here confirm that the target antigen selection critically impacts antibody efficacy in vitro.^66^ For example, Fc-mediated tumor cell killing was notoriously weak using unmodified wildtype human IgG1 antibodies against CD19,^16^ but could be significantly improved by respective Fc engineering.^38^ Consequently, the clinically available CD19 antibody TAFA is engineered for improved FcR binding compared to wildtype IgG1 and also demonstrated preclinical activity in depleting B cells against B-ALL in PDX studies^18,30^ and in non-human primate models.^16^ Further enhancement of ADCP in vitro and therapeutic activity in vivo was observed in combination with CD47 blockade,^30^ which – at least in vitro – showed limited efficacy for a non-engineered CD19 antibody containing the TAFA variable regions. Furthermore, a CD89-directed bispecific antibody^67^ or isotype switching of CD19 antibodies to IgA did not recruit PMN for ADCC – the latter not even in the presence of CD47 blockade. These results with myeloid effector cells contrast to observations with T cells, where CD19-directed bispecific antibodies or CD19-directed CARs induce potent killing of leukemia and lymphoma cells.^64,68^ These differences may be related to the fundamentally different tumor cell killing mechanisms of T cells compared to myeloid cells: while T cells kill via FAS/FAS ligand interactions^69^ or induce membrane pores into tumor cells,^70^ macrophages kill tumor cells by phagocytosis^12^ and PMN by “frustrated” phagocytosis, called trogoptosis.^71^ While mechanisms of sensitivity and resistance against tumor cell killing by T and NK cells or macrophages begin to emerge,^72,73,74^ less is known about molecular mechanisms affecting the efficacy of trogoptosis.^75^

In line with results from lymphoma studies,^14,17^ CD20 antibodies effectively triggered ADCP and ADCC also against CD20 expressing B-ALL cells, which was further enhanced by CD47 blockade. Effective ADCP by macrophages was observed with the glyco-engineered OBZ, but also with the non-engineered RTX and with IgA2 variants of both antibodies. As expected from experiments with lymphoma cells,^39^ ADCC by PMN was only observed with anti-CD20 IgA2 variants, but not with their clinically available IgG1 versions – irrespective of CD47 blockade. This difference between IgG1 and IgA2 isotypes in triggering ADCC by PMN is probably explained by differences of antibody binding to FcR, leading to more potent downstream signaling of CD89 compared to Fcγ receptors in PMN.^24^ An obvious limitation of all CD20 antibodies in the treatment of B-ALL is the restricted expression of CD20 on certain subtypes, which is less obvious from mRNA studies compared to classical immunological classifications where early pro-B-ALL samples like NALM-6 or SEM are typically CD20 negative. Nevertheless, RTX is nowadays widely used in the treatment of adult B-ALL patients.^3^ Approaches to increase the efficacy of CD20 directed antibody therapy – e.g. by switching to OBZ, to IgA2 variants, or to combinations with CD47 blockade – require further studies.

The CD38 antibody DARA continues to change the therapeutic landscape for plasma cell dyscrasias,^76^ and CD38 expression is also found on many B- and T-ALL leukemia cells. Preclinical in vitro and in vivo studies have demonstrated that CD38 antibodies like DARA and isatuximab could also have therapeutic potential for the treatment of ALL patients.^57^ Recently published results from a phase II study showed promising results for DARA in T-ALL patients, while results in B-ALL were less impressive.^77^ Our results presented here demonstrate that myeloid effector cell recruitment by DARA could be improved by switching isotypes from IgG1 to IgA2, and by combining CD38 antibodies with CD47 blockade. These effects were observed for both macrophages and PMN. The in vivo study confirmed FcαRI (CD89)-dependent efficacy of anti-CD38 IgA2, which was significantly enhanced by a combination with a SIRPα-Fc fusion protein. Contrary to the in vitro data, however, CD47 blockade alone was also highly effective in vivo, which has also been observed in other ALL models^29^ and may reflect CD47 dependent tumor cell engraftment.^78^ While clinical studies with the CD47 antibody magrolimab were discontinued,^32^ clinical studies with SIRPα antibodies and a SIRPα-Fc fusion protein are on-going.^79^ Importantly, an unbiased screening approach found combined CD47 and CD38 targeting particularly effective in killing malignant B cells,^80^ and clinical studies with CD47xCD19 and CD47xCD38 bispecific antibodies are ongoing in lymphoma and myeloma patients, respectively.^81,82^ Depending on their toxicity and efficacy in these other indications, they can also be considered for testing in ALL therapy. An additional option to interfere with CD47/SIRPα interactions could be small molecule QPCTL inhibitors,^63^ which were demonstrated to enhance ADCP and ADCC^34,35^ and to favorably modify the tumor microenvironment.^33^

IgA antibodies induce potent tumor cell killing against several target antigens in vitro and in vivo.^83^ However, their clinical development faces several challenges. Nevertheless, many biotech companies have or had ongoing IgA programs, but the first IgA antibody still has to enter clinical trials. For the concept of myeloid checkpoint blockade in antibody-based immunotherapy, IgA antibodies appear to be an ideal combination partner, since IgA effectively activates both macrophages and PMN. Potentially, ALL could become a model disease again when IgA variants of CD38 antibodies would be developed together with CD47/SIRPα interference.

## Acknowledgements

This research study was funded by the German Research Foundation (DFG; Clinical Research Unit “CATCH ALL” KFO 5010/1 to AH, LB, MB, FS, MP, CDB, DMS and TV. DFG INST 186/1465-1 to KS and DFG INST 186/1230-1 FUGG supported the Q Exactive/Ultimate 3000 LC-MS). ML and TV were supported by intramural funding by the Medical Faculty of the CAU. ICP was supported by Villa Joep Foundation. MJ received funding by the Dutch Cancer Society (KWF-PPS2023-15556). JL was supported by Oncode Accelerator, a Dutch National Growth Fund project under grant number NGFOP2201. We gratefully acknowledge M.Sc. Alexander Jochimsen for help in producing anti-CD38 IgA2 and Vanessa Möller for her contribution to the cross-validation experiments. We thank Dr. Arturo Macarrón Palacios from GenCC GmbH as well as Dr. Pipob Suwanchaikasem and Dr. Christine Joy I. Bulaon from Baiya Phytopharm for peptide mapping and glyco-profiling of anti-CD38 IgA2.

## Authors’ contribution

MK, ML, NB, IP, DW, SK, TR, SB, MB, AAW and KS performed in vitro experiments; MJ, IP and JL were responsible for the in vivo studies. AH, SB, LB and OV performed data analyses. MK, ML, RB, FS, CB, MP, DMS, JL and TV designed the study. MK, ML, RB and TV wrote the manuscript, which was critically reviewed and approved by all authors before submission.

## List of abbreviations

ADCC: antibody-dependent cell-mediated cytotoxicity
ADCP: antibody-dependent cellular phagocytosis
B-ALL B: cell precursor acute lymphoblastic leukemia
BLI: bioluminescence imaging
CAR: chimeric antigen receptor
CD: cluster of differentiation
CPM (transcriptomics): counts per million
CPM (radioactivity-based readout): counts per minute
CTX: cetuximab
DARA: daratumumab
EGA: European Genome-Phenome Archive
FcR: fragment crystallizable receptor
FITC: fluorescein isothiocyanate
G(M)-CSF: granulocyte (macrophage) colony-stimulating factor
HRP: horseradish peroxidase
i.p.: intraperitoneal
ITAM: immunoreceptor tyrosine-based activation motif
ITIM: immunoreceptor tyrosine-based inhibition motif
LC: liquid chromatography
LUC: firefly luciferase
M0: non-polarized macrophages
MS: mass spectrometry
MDS: myelodysplastic syndrome
MDSC: myeloid-derived suppressor cells
NXG: non-obese diabetic xenograft gamma strain
OBZ: obinutuzumab
PBMC: peripheral blood mononuclear cells
Peg: pegylated (conjugated with polyethylene glycol)
pGlu: pyroglutamic acid
PMN: polymorphonuclear leukocytes
PNGase: F eptide-N-Glycosidase F
QPCTL: glutaminyl-peptide cyclotransferase-like protein
ROI: red objects per image
RTX: rituximab
SABC: specific antibody binding capacity
SIRPα: signal regulatory protein α
TAFA: tafasitamab
UMAP: uniform manifold approximation and projection

## References

1. Gökbuget N, Boissel N, Chiaretti S, Dombret H, Doubek M, Fielding A, Foà R, Giebel S, Hoelzer D, Hunault M, Marks DI, Martinelli G, Ottmann O, Rijneveld A, Rousselot P, Ribera J, Bassan R. Management of ALL in adults: 2024 ELN recommendations from a European expert panel. Blood. 2024;143(19):1903–1930. doi:10.1182/blood.2023023568

2. Jabbour E, Short NJ, Jain N, Haddad FG, Welch MA, Ravandi F, Kantarjian H. The evolution of acute lymphoblastic leukemia research and therapy at MD Anderson over four decades. J Hematol Oncol. 2023;16(1):22. doi:10.1186/s13045-023-01409-5

3. Badar T, Luger SM, Litzow MR. Incorporation of immunotherapy into frontline treatment for adults with B-cell precursor acute lymphoblastic leukemia. Blood. 2025;145(14):1475–1484. doi:10.1182/blood.2023022921

4. June CH, Sadelain M. Chimeric antigen receptor therapy. New Engl J Med. 2018;379(1):64–73. doi:10.1056/NEJMra1706169

5. Gupta S, Rau RE, Kairalla JA, Rabin KR, Wang C, Angiolillo AL, Alexander S, Carroll AJ, Conway S, Gore L, Kirsch I, Kubaney HR, Li AM, McNeer JL, Militano O, Miller TP, Moyer Y, O’Brien MM, Okada M, Reshmi SC, Shago M, Wagner E, Winick N, Wood BL, Haworth-Wright T, Zaman F, Zugmaier G, Zupanec S, Devidas M, Hunger SP, Teachey DT, Raetz EA, Loh ML. Blinatumomab in standard risk pediatric B-acute lymphoblastic leukemia. N Engl J Med. 2025;392(9):875–891. doi:10.1056/NEJMoa2411680

6. Maury S, Chevret S, Thomas X, Heim D, Leguay T, Huguet F, Chevallier P, Hunault M, Boissel N, Escoffre-Barbe M, Hess U, Vey N, Pignon JM, Braun T, Marolleau JP, Cahn JY, Chalandon Y, Lhéritier V, Beldjord K, Béné MC, Ifrah N, Dombret H. Rituximab in B-lineage adult acute lymphoblastic leukemia. N Engl J Med. 2016;375(11):1044–1053. doi:10.1056/NEJMoa1605085

7. Kantarjian HM, DeAngelo DJ, Stelljes M, Martinelli G, Liedtke M, Stock W, Gökbuget N, O’Brien S, Wang K, Wang T, Paccagnella ML, Sleight B, Vandendries E, Advani AS. Inotuzumab ozogamicin versus standard therapy for acute lymphoblastic leukemia. N Engl J Med. 2016;375(8):740–753. doi:10.1056/NEJMoa1509277

8. Gentles AJ, Newman AM, Liu CL, Bratman SV, Feng W, Kim D, Nair VS, Xu Y, Khuong A, Hoang CD, Diehn M, West RB, Plevritis SK, Alizadeh AA. The prognostic landscape of genes and infiltrating immune cells across human cancers. Nat Med. 2015;21(8):938–945. doi:10.1038/nm.3909

9. Vlerken-Ysla L van, Tyurina YY, Kagan VE, Gabrilovich DI. Functional states of myeloid cells in cancer. Cancer Cell. 2023;41(3):490–504. doi:10.1016/j.ccell.2023.02.009

10. Witkowski MT, Dolgalev I, Evensen NA, Ma C, Chambers T, Roberts KG, Sreeram S, Dai Y, Tikhonova AN, Lasry A, Qu C, Pei D, Cheng C, Robbins GA, Pierro J, Selvaraj S, Mezzano V, Daves M, Lupo PJ, Scheurer ME, Loomis CA, Mullighan CG, Chen W, Rabin KR, Tsirigos A, Carroll WL, Aifantis I. Extensive remodeling of the immune microenvironment in B cell acute lymphoblastic leukemia. Cancer Cell. 2020;37(6):867–882.e12. doi:10.1016/j.ccell.2020.04.015

11. Mantovani A, Allavena P, Marchesi F, Garlanda C. Macrophages as tools and targets in cancer therapy. Nat Rev Drug Discov. 2022;21(11):799–820. doi:10.1038/s41573-022-00520-5

12. Van Wagoner CM, Rivera-Escalera F, Jaimes-Delgadillo NC, Chu CC, Zent CS, Elliott MR. Antibody-mediated phagocytosis in cancer immunotherapy. Immunol Rev. 2023;319(1):128–141. doi:10.1111/imr.13265

13. Uchida J, Hamaguchi Y, Oliver JA, Ravetch JV, Poe JC, Haas KM, Tedder TF. The innate mononuclear phagocyte network depletes B lymphocytes through Fc receptor– dependent mechanisms during anti-CD20 antibody immunotherapy. J Exp Med. 2004;199(12):1659–1669. doi:10.1084/jem.20040119

14. Minard-Colin V, Xiu Y, Poe JC, Horikawa M, Magro CM, Hamaguchi Y, Haas KM, Tedder TF. Lymphoma depletion during CD20 immunotherapy in mice is mediated by macrophage FcγRI, FcγRIII, and FcγRIV. Blood. 2008;112(4):1205–1213. doi:10.1182/blood-2008-01-135160

15. Grandjean CL, Montalvao F, Celli S, Michonneau D, Breart B, Garcia Z, Perro M, Freytag O, Gerdes CA, Bousso P. Intravital imaging reveals improved Kupffer cell-mediated phagocytosis as a mode of action of glycoengineered anti-CD20 antibodies. Sci Rep. 2016;6:34382. doi:10.1038/srep34382

16. Horton HM, Bernett MJ, Pong E, Peipp M, Karki S, Chu SY, Richards JO, Vostiar I, Joyce PF, Repp R, Desjarlais JR, Zhukovsky EA. Potent in vitro and in vivo activity of an Fc-engineered anti-CD19 monoclonal antibody against lymphoma and leukemia. Cancer Res. 2008;68(19):8049–8057. doi:10.1158/0008-5472.CAN-08-2268

17. Herter S, Herting F, Mundigl O, Waldhauer I, Weinzierl T, Fauti T, Muth G, Ziegler-Landesberger D, Van Puijenbroek E, Lang S, Duong MN, Reslan L, Gerdes CA, Friess T, Baer U, Burtscher H, Weidner M, Dumontet C, Umana P, Niederfellner G, Bacac M, Klein C. Preclinical activity of the type II CD20 antibody GA101 (obinutuzumab) compared with rituximab and ofatumumab in vitro and in xenograft models. Mol Cancer Ther. 2013;12(10):2031–2042. doi:10.1158/1535-7163.MCT-12-1182

18. Schewe DM, Alsadeq A, Sattler C, Lenk L, Vogiatzi F, Cario G, Vieth S, Valerius T, Rosskopf S, Meyersieck F, Alten J, Schrappe M, Gramatzki M, Peipp M, Kellner C. An Fc-engineered CD19 antibody eradicates MRD in patient-derived MLL-rearranged acute lymphoblastic leukemia xenografts. Blood. 2017;130(13):1543–1552. doi:10.1182/blood-2017-01-764316

19. Vogiatzi F, Winterberg D, Lenk L, Buchmann S, Cario G, Schrappe M, Peipp M, Richter-Pechanska P, Kulozik AE, Lentes J, Bergmann AK, Valerius T, Frielitz FS, Kellner C, Schewe DM. Daratumumab eradicates minimal residual disease in a preclinical model of pediatric T-cell acute lymphoblastic leukemia. Blood. 2019;134(8):713–716. doi:10.1182/blood.2019000904

20. Derer S, Glorius P, Schlaeth M, Lohse S, Klausz K, Muchhal U, Desjarlais JR, Humpe A, Valerius T, Peipp M. Increasing FcγRIIa affinity of an FcγRIII-optimized anti-EGFR antibody restores neutrophil-mediated cytotoxicity. MAbs. 2014;6:409–421. 10.4161/mabs.27457

21. Treffers LW, van Houdt M, Bruggeman CW, Heineke MH, Zhao XW, van der Heijden J, Nagelkerke SQ, Verkuijlen PJJH, Geissler J, Lissenberg-Thunnissen S, Valerius T, Peipp M, Franke K, van Bruggen R, Kuijpers TW, van Egmond M, Vidarsson G, Matlung HL, van den Berg TK. FcγRIIIb restricts antibody-dependent destruction of cancer cells by human neutrophils. Front Immunol. 2018;9:3124. doi:10.3389/fimmu.2018.03124

22. Boross P, Lohse S, Nederend M, Jansen JHM, van Tetering G, Dechant M, Peipp M, Royle L, Liew LP, Boon L, van Rooijen N, Bleeker WK, Parren PWHI, van de Winkel JGJ, Valerius T, Leusen JHW. IgA EGFR antibodies mediate tumour killing in vivo. EMBO Mol Med. 2013;5(8):1213–1226. doi:10.1002/emmm.201201929

23. Lohse S, Meyer S, Meulenbroek LAPM, Jansen JHM, Nederend M, Kretschmer A, Klausz K, Möginger U, Derer S, Rösner T, Kellner C, Schewe D, Sondermann P, Tiwari S, Kolarich D, Peipp M, Leusen JHW, Valerius T. An anti-EGFR IgA that displays improved pharmacokinetics and myeloid effector cell engagement in vivo. Cancer Res. 2016;76(2):403–417. doi:10.1158/0008-5472.CAN-15-1232

24. Brandsma AM, Bondza S, Evers M, Koutstaal R, Nederend M, Jansen JHM, Rösner T, Valerius T, Leusen JHW, Ten Broeke T. Potent Fc receptor signaling by IgA leads to superior killing of cancer cells by neutrophils compared to IgG. Front Immunol. 2019;10:704. doi:10.3389/fimmu.2019.00704

25. Treffers LW, Ten Broeke T, Rösner T, Jansen JHM, van Houdt M, Kahle S, Schornagel K, Verkuijlen PJJH, Prins JM, Franke K, Kuijpers TW, van den Berg TK, Valerius T, Leusen JHW, Matlung HL. IgA-mediated killing of tumor cells by neutrophils is enhanced by CD47-SIRPα checkpoint inhibition. Cancer Immunol Res. 2020;8(1):120–130. doi:10.1158/2326-6066.CIR-19-0144

26. Chan C, Stip M, Nederend M, Jansen M, Passchier E, van den Ham F, Wienke J, van Tetering G, Leusen J. Enhancing IgA-mediated neutrophil cytotoxicity against neuroblastoma by CD47 blockade. J Immunother Cancer. 2024;12(5):e008478. doi:10.1136/jitc-2023-008478

27. Barclay AN, van den Berg TK. The interaction between signal regulatory protein alpha (SIRPα) and CD47: structure, function, and therapeutic target. Annu Rev Immunol. 2014;32(1):25–50. doi:10.1146/annurev-immunol-032713-120142

28. Maute R, Xu J, Weissman IL. CD47–SIRPα-targeted therapeutics: status and prospects. Immunooncol Technol. 2022;13:100070. doi:10.1016/j.iotech.2022.100070

29. Müller K, Vogiatzi F, Winterberg D, Rösner T, Lenk L, Bastian L, Gehlert CL, Autenrieb MP, Brüggemann M, Cario G, Schrappe M, Kulozik AE, Eckert C, Bergmann AK, Bornhauser B, Bourquin JP, Valerius T, Peipp M, Kellner C, Schewe DM. Combining daratumumab with CD47 blockade prolongs survival in preclinical models of pediatric T-ALL. Blood. 2022;140(1):45–57. doi:10.1182/blood.2021014485

30. Schewe DM, Vogiatzi F, Münnich IA, Zeller T, Windisch R, Wichmann C, Müller K, Bhat H, Felix E, Mougiakakos D, Bruns H, Lenk L, Valerius T, Humpe A, Peipp M, Kellner C. Enhanced potency of immunotherapy against B-cell precursor acute lymphoblastic leukemia by combination of an Fc-engineered CD19 antibody and CD47 blockade. Hemasphere. 2024;8(2):e48. doi:10.1002/hem3.48

31. Advani R, Flinn I, Popplewell L, Forero A, Bartlett NL, Ghosh N, Kline J, Roschewski M, LaCasce A, Collins GP, Tran T, Lynn J, Chen JY, Volkmer JP, Agoram B, Huang J, Majeti R, Weissman IL, Takimoto CH, Chao MP, Smith SM. CD47 blockade by Hu5F9-G4 and rituximab in Non-Hodgkin’s lymphoma. N Engl J Med. 2018;379(18):1711–1721. doi:10.1056/NEJMoa1807315

32. Gilead Sciences. Gilead statement on discontinuation of phase 3 ENHANCE-3 study in AML. www.gilead.com. February 7, 2024. Accessed December 2, 2024. https://www.gilead.com/company/company-statements/2024/gilead-statement-on-discontinuation-of-phase-3-enhance-3-study-in-aml

33. Barreira da Silva R, Leitao RM, Pechuan-Jorge X, Werneke S, Oeh J, Javinal V, Wang Y, Phung W, Everett C, Nonomiya J, Arnott D, Lu C, Hsiao YC, Koerber JT, Hötzel I, Ziai J, Modrusan Z, Pillow TH, Roose-Girma M, Schartner JM, Merchant M, Rutz S, Eidenschenk C, Mellman I, Albert ML. Loss of the intracellular enzyme QPCTL limits chemokine function and reshapes myeloid infiltration to augment tumor immunity. Nat Immunol. 2022;23(4):568–580. doi:10.1038/s41590-022-01153-x

34. Logtenberg MEW, Jansen JHM, Raaben M, Toebes M, Franke K, Brandsma AM, Matlung HL, Fauster A, Gomez-Eerland R, Bakker NAM, van der Schot S, Marijt KA, Verdoes M, Haanen JBAG, van den Berg JH, Neefjes J, van den Berg TK, Brummelkamp TR, Leusen JHW, Scheeren FA, Schumacher TN. Glutaminyl cyclase is an enzymatic modifier of the CD47- IRPα axis and a target for cancer immunotherapy. Nat Med. 2019;25(4):612–619. doi:10.1038/s41591-019-0356-z

35. Wu Z, Weng L, Zhang T, Tian H, Fang L, Teng H, Zhang W, Gao J, Hao Y, Li Y, Zhou H, Wang P. Identification of glutaminyl cyclase isoenzyme isoQC as a regulator of SIRPα-CD47 axis. Cell Res. 2019;29(6):502–505. doi:10.1038/s41422-019-0177-0

36. Beder T, Hansen BT, Hartmann AM, Zimmermann J, Amelunxen E, Wolgast N, Walter W, Zaliova M, Antić Ž, Chouvarine P, Bartsch L, Barz MJ, Bultmann M, Horns J, Bendig S, Kässens J, Kaleta C, Cario G, Schrappe M, Neumann M, Gökbuget N, Bergmann AK, Trka J, Haferlach C, Brüggemann M, Baldus CD, Bastian L. The gene expression classifier ALLCatchR identifies B-cell precursor ALL subtypes and underlying developmental trajectories across age. Hemasphere. 2023;7(9):e939. doi:10.1097/HS9.0000000000000939

37. Baumann N, Arndt C, Petersen J, Lustig M, Rösner T, Klausz K, Kellner C, Bultmann M, Bastian L, Vogiatzi F, Leusen JHW, Burger R, Schewe DM, Peipp M, Valerius T. Myeloid checkpoint blockade improves killing of T-acute lymphoblastic leukemia cells by an IgA2 variant of daratumumab. Front Immunol. 2022;13:949140. doi:10.3389/fimmu.2022.949140

38. Gehlert CL, Rahmati P, Boje AS, Winterberg D, Krohn S, Theocharis T, Cappuzzello E, Lux A, Nimmerjahn F, Ludwig RJ, Lustig M, Rösner T, Valerius T, Schewe DM, Kellner C, Klausz K, Peipp M. Dual Fc optimization to increase the cytotoxic activity of a CD19-targeting antibody. Front Immunol. 2022;13:957874. doi:10.3389/fimmu.2022.957874

39. Evers M, Rösner T, Dünkel A, Jansen JHM, Baumann N, Ten Broeke T, Nederend M, Eichholz K, Klausz K, Reiding K, Schewe DM, Kellner C, Peipp M, Leusen JHW, Valerius T. The selection of variable regions affects effector mechanisms of IgA antibodies against CD20. Blood Adv. 2021;5(19):3807–3820. doi:10.1182/bloodadvances.2021004598

40. Chernyavska M, Hermans CKJC, Chan C, Baumann N, Rösner T, Leusen JHW, Valerius T, Verdurmen WPR. Evaluation of immunotherapies improving macrophage anti-tumor response using a microfluidic model. Organs-on-a-Chip. 2022;4:100019. doi:10.1016/j.ooc.2022.100019

41. Peipp M, Lammerts van Bueren JJ, Schneider-Merck T, Bleeker WWK, Dechant M, Beyer T, Repp R, van Berkel PHC, Vink T, van de Winkel JGJ, Parren PWHI, Valerius T. Antibody fucosylation differentially impacts cytotoxicity mediated by NK and PMN effector cells. Blood. 2008;112(6):2390–2399. doi:10.1182/blood-2008-03-144600

42. Lustig M, Hahn C, Leangen Herigstad M, Andersen JT, Leusen JHW, Burger R, Valerius T. Sialylation inhibition improves macrophage mediated tumor cell phagocytosis of breast cancer cells triggered by therapeutic antibodies of different isotypes. Front Oncol. 2024;14. doi:10.3389/fonc.2024.1488668

43. Stip MC, Jansen JHM, Nederend M, Tsioumpekou M, Evers M, Olofsen PA, Meyer-Wentrup F, Leusen JHW. Characterization of human Fc alpha receptor transgenic mice: comparison of CD89 expression and antibody-dependent tumor killing between mouse strains. Cancer Immunol Immunother. 2023;72(9):3063–3077. doi:10.1007/s00262-023-03478-4

44. Baumann N, Rösner T, Jansen JHM, Chan C, Marie Eichholz K, Klausz K, Winterberg D, Müller K, Humpe A, Burger R, Peipp M, Schewe DM, Kellner C, Leusen JHW, Valerius T. Enhancement of epidermal growth factor receptor antibody tumor immunotherapy by glutaminyl cyclase inhibition to interfere with CD47/signal regulatory protein alpha interactions. Cancer Sci. 2021;112(8):3029–3040. doi:10.1111/cas.14999

45. Kassambara A. ggpubr: “ggplot2” Based Publication Ready Plots_. R package version 0.6.0, <https://CRAN.R-project.org/package=ggpubr>. Published online February 10, 2023. Accessed January 7, 2025. https://cran.r-project.org/web/packages/ggpubr/index.html

46. Bene MC, Castoldi G, Knapp W, Ludwig WD, Matutes E, Orfao A, van’t Veer MB. Proposals for the immunological classification of acute leukemias. European Group for the Immunological Characterization of Leukemias (EGIL). Leukemia. 1995;9(10):1783–1786.

47. Rolink A, Melchers F. Generation and regeneration of cells of the B-lymphocyte lineage. Curr Opin Immunol. 1993;5(2):207–217. doi:10.1016/0952-7915(93)90006-E

48. LeBien TW, Tedder TF. B lymphocytes: how they develop and function. Blood. 2008;112(5):1570–1580. doi:10.1182/blood-2008-02-078071

49. Passet M, Kim R, Clappier E. Genetic subtypes of B-cell acute lymphoblastic leukemia in adults. Blood. 2025;145(14):1451–1463. doi:10.1182/blood.2023022919

50. Derer S, Lohse S, Valerius T. EGFR expression levels affect the mode of action of EGFR-targeting monoclonal antibodies. Oncoimmunology. 2013;2(5):e24052. doi:10.4161/onci.24052

51. Cleary KLS, Chan HTC, James S, Glennie MJ, Cragg MS. Antibody distance from the cell membrane regulates antibody effector mechanisms. J Immunol. 2017;198(10):3999–4011. doi:10.4049/jimmunol.1601473

52. Singhal S, Rao AS, Stadanlick J, Bruns K, Sullivan NT, Bermudez A, Honig-Frand A, Krouse R, Arambepola S, Guo E, Moon EK, Georgiou G, Valerius T, Albelda SM, Eruslanov EB. Human tumor–associated macrophages and neutrophils regulate antitumor antibody efficacy through lethal and sublethal trogocytosis. Cancer Res. 2024;84(7):1029. doi:10.1158/0008-5472.CAN-23-2135

53. Kolde R. pheatmap: Pretty Heatmaps. R package version 1.0.12, <https://CRAN.R-project.org/package=pheatmap>. Published online 2019. doi:10.32614/CRAN.package.pheatmap

54. Brivio E, Bautista F, Zwaan CM. Naked antibodies and antibody-drug conjugates: targeted therapy for childhood acute lymphoblastic leukemia. Haematologica. 2024;109(6):1700–1712. doi:10.3324/haematol.2023.283815

55. van den Berg TK, Valerius T. Myeloid immune-checkpoint inhibition enters the clinical stage. Nat Rev Clin Oncol. 2019;16(5):275–276. doi:10.1038/s41571-018-0155-3

56. Maoz A, Weiskopf K. Phagocytic cooperativity by tumour macrophages. Nat Biomed Eng. 2023;7(9):1057–1059. doi:10.1038/s41551-023-01088-0

57. Senapati J, Jabbour E. Dar-ting at CD38 in T-cell leukemias. Blood. 2024;144(21):2162–2164. doi:10.1182/blood.2024026118

58. Stockmeyer B, Dechant M, van Egmond M, Tutt AL, Sundarapandiyan K, Graziano RF, Repp R, Kalden JR, Gramatzki M, Glennie MJ, van de Winkel JG, Valerius T. Triggering Fc alpha-receptor I (CD89) recruits neutrophils as effector cells for CD20-directed antibody therapy. J Immunol. 2000;165(10):5954–5961. doi:10.4049/jimmunol.165.10.5954

59. Montoyo HP, Vaccaro C, Hafner M, Ober RJ, Mueller W, Ward ES. Conditional deletion of the MHC class I-related receptor FcRn reveals the sites of IgG homeostasis in mice. Proc Natl Acad Sci USA. 2009;106(8):2788–2793. doi:10.1073/pnas.0810796106

60. Huang KF, Liu YL, Cheng WJ, Ko TP, Wang AHJ. Crystal structures of human glutaminyl cyclase, an enzyme responsible for protein N-terminal pyroglutamate formation. Proc Natl Acad Sci U S A. 2005;102(37):13117–13122. doi:10.1073/pnas.0504184102

61. Hatherley D, Graham SC, Turner J, Harlos K, Stuart DI, Barclay AN. Paired receptor specificity explained by structures of signal regulatory proteins alone and complexed with CD47. Mol Cell. 2008;31(2):266–277. doi:10.1016/j.molcel.2008.05.026

62. Tam SH, McCarthy SG, Armstrong AA, Somani S, Wu SJ, Liu X, Gervais A, Ernst R, Saro D, Decker R, Luo J, Gilliland GL, Chiu ML, Scallon BJ. Functional, Biophysical, and Structural Characterization of Human IgG1 and IgG4 Fc Variants with Ablated Immune Functionality. Antibodies (Basel). 2017;6(3):12. doi:10.3390/antib6030012

63. Yu L, Sun Y, Xie L, Tan X, Wang P, Xu S. Targeting QPCTL: An emerging therapeutic opportunity. J Med Chem. 2025;68(2):929–943. doi:10.1021/acs.jmedchem.4c02247

64. Kantarjian H, Stein A, Gökbuget N, Fielding AK, Schuh AC, Ribera JM, Wei A, Dombret H, Foà R, Bassan R, Arslan Ö, Sanz MA, Bergeron J, Demirkan F, Lech-Maranda E, Rambaldi A, Thomas X, Horst HA, Brüggemann M, Klapper W, Wood BL, Fleishman A, Nagorsen D, Holland C, Zimmerman Z, Topp MS. Blinatumomab versus chemotherapy for advanced acute lymphoblastic leukemia. N Engl J Med. 2017;376(9):836–847. doi:10.1056/NEJMoa1609783

65. Kegyes D, Ghiaur G, Bancos A, Tomuleasa C, Gale RP. Immune therapies of B-cell acute lymphoblastic leukaemia in children and adults. Crit Rev Oncol Hematol. 2024;196:104317. doi:10.1016/j.critrevonc.2024.104317

66. Würflein D, Dechant M, Stockmeyer B, Tutt AL, Hu P, Repp R, Kalden JR, van de Winkel JG, Epstein AL, Valerius T, Glennie M, Gramatzki M. Evaluating antibodies for their capacity to induce cell-mediated lysis of malignant B cells. Cancer Res. 1998;58(14):3051–3058.

67. Tiroch K, Stockmeyer B, Frank C, Valerius T. Intracellular domains of target antigens influence their capacity to trigger antibody-dependent cell-mediated cytotoxicity. J Immunol. 2002;168(7):3275–3282. doi:10.4049/jimmunol.168.7.3275

68. Larson RC, Maus MV. Recent advances and discoveries in the mechanisms and functions of CAR T cells. Nat Rev Cancer. 2021;21(3):145–161. doi:10.1038/s41568-020-00323-z

69. Loves R, Grunebaum E. FAS signalling pathway is crucial for CAR T cell persistence. Nat Rev Immunol. 2024;24(6):380–380. doi:10.1038/s41577-024-01038-0

70. Arenas EJ, Martínez-Sabadell A, Rius Ruiz I, Román Alonso M, Escorihuela M, Luque A, Fajardo CA, Gros A, Klein C, Arribas J. Acquired cancer cell resistance to T cell bispecific antibodies and CAR T targeting HER2 through JAK2 down-modulation. Nat Commun. 2021;12(1):1237. doi:10.1038/s41467-021-21445-4

71. Matlung HL, Babes L, Zhao XW, Houdt M van, Treffers LW, Rees DJ van, Franke K, Schornagel K, Verkuijlen P, Janssen H, Halonen P, Lieftink C, Beijersbergen RL, Leusen JHW, Boelens JJ, Kuhnle I, Bosch J van der WT, Seeger K, Rutella S, Pagliara D, Matozaki T, Suzuki E, Oordt CWM van der H van, Bruggen R van, Roos D, Lier RAW van, Kuijpers TW, Kubes P, Berg TK van den. Neutrophils kill antibody-opsonized cancer cells by trogoptosis. Cell Rep. 2018;23(13):3946–3959.e6. doi:10.1016/j.celrep.2018.05.082

72. Freeman AJ, Vervoort SJ, Ramsbottom KM, Kelly MJ, Michie J, Pijpers L, Johnstone RW, Kearney CJ, Oliaro J. Natural Killer Cells Suppress T Cell-Associated Tumor Immune Evasion. Cell Rep. 2019;28(11):2784–2794.e5. doi:10.1016/j.celrep.2019.08.017

73. Cantoni C, Wurzer H, Thomas C, Vitale M. Escape of tumor cells from the NK cell cytotoxic activity. J Leukoc Biol. 2020;108(4):1339–1360. doi:10.1002/JLB.2MR0820-652R

74. Yang Y long, Yang F, Huang Z qing, Li Y yuan, Shi H yuan, Sun Q, Ma Y, Wang Y, Zhang Y, Yang S, Zhao G ren, Xu F hua. T cells, NK cells, and tumor-associated macrophages in cancer immunotherapy and the current state of the art of drug delivery systems. Front Immunol. 2023;14:1199173. doi:10.3389/fimmu.2023.1199173

75. Bouti P, Klein BJAM, Verkuijlen PJH, Schornagel K, Van Alphen FPJ, Taris KKH, Van Den Biggelaar M, Hoogendijk AJ, Van Bruggen R, Kuijpers TW, Matlung HL. SKAP2 acts downstream of CD11b/CD18 and regulates neutrophil effector function. Front Immunol. 2024;15:1344761. doi:10.3389/fimmu.2024.1344761

76. Maura F, Bergsagel PL. Targeting the Tumor and the Immune System in Smoldering Multiple Myeloma. New England Journal of Medicine. 2025;392(18):1858–1860. doi:10.1056/NEJMe2504273

77. Bhatla T, Hogan LE, Teachey DT, Bautista F, Moppett J, Velasco Puyó P, Micalizzi C, Rossig C, Shukla N, Gilad G, Locatelli F, Baruchel A, Zwaan CM, Bezler NS, Rubio-San-Simón A, Taussig DC, Raetz EA, Mao ZJ, Wood BL, Alvarez Arias D, Krevvata M, Nnane I, Bandyopadhyay N, Lopez Solano L, Dennis RM, Carson R, Vora A. Daratumumab in pediatric relapsed/refractory acute lymphoblastic leukemia or lymphoblastic lymphoma: the DELPHINUS study. Blood. 2024;144(21):2237–2247. doi:10.1182/blood.2024024493

78. Chao MP, Weissman IL, Majeti R. The CD47–SIRPα pathway in cancer immune evasion and potential therapeutic implications. Curr Opin Immunol. 2012;24(2):225–232. doi:10.1016/j.coi.2012.01.010

79. Yang H, Xun Y, You H. The landscape overview of CD47-based immunotherapy for hematological malignancies. Biomark Res. 2023;11(1):15. doi:10.1186/s40364-023-00456-x

80. Ribeiro J, Pagès-Geli C, Meglan A, Velarde J, Blandin J, Vaccaro K, Wienclaw T, Fernández-Guzmán P, Hahn CK, Crespo M, Weiskopf K. Unbiased discovery of antibody therapies that stimulate macrophage-mediated destruction of B-cell lymphoma. bioRxiv [Preprint]. Published online November 15, 2024:2024.11.13.623229. doi:10.1101/2024.11.13.623229

81. Hawkes EA, Lewis KL, Doo NW, Patil SS, Miskin HP, Sportelli P, Kolibaba KS, Normant E, Turpuseema T, Cheah CY. First-in-human (FIH) study of the fully-human kappa-lambda CD19/CD47 bispecific antibody TG-1801 in patients with B-cell lymphoma. Hematol Oncol. 2023;41(S2):579–580. doi:10.1002/hon.3164_434

82. Grandclément C, Estoppey C, Dheilly E, Panagopoulou M, Monney T, Dreyfus C, Loyau J, Labanca V, Drake A, De Angelis S, Rubod A, Frei J, Caro LN, Blein S, Martini E, Chimen M, Matthes T, Kaya Z, Edwards CM, Edwards JR, Menoret E, Kervoelen C, Pellat-Deceunynck C, Moreau P, Mbow ML, Srivastava A, Dyson MR, Zhukovsky EA, Perro M, Sammicheli S. Development of ISB 1442, a CD38 and CD47 bispecific biparatopic antibody innate cell modulator for the treatment of multiple myeloma. Nat Commun. 2024;15(1):2054. doi:10.1038/s41467-024-46310-y

83. van Tetering G, Evers M, Chan C, Stip M, Leusen J. Fc engineering strategies to advance IgA antibodies as therapeutic agents. Antibodies. 2020;9(4):70. doi:10.3390/antib9040070

