## Supplement for "Myeloid cell-mediated killing of B-ALL by CD38 and CD20 IgA antibody variants is enhanced by CD47/SIRPα interference"

### Supplemental Materials and Methods

#### RNA profiling of patient samples and cell lines

For whole transcriptome sequencing, total RNA from each cell line was extracted using TRIzol (Thermo Fisher Scientific; cat. no. 15596026) and its integrity was verified by agarose-formaldehyde gel electrophoresis as described.[1] Library preparation was performed with 100 ng RNA input using the Illumina mRNA Stranded Kit (Illumina®; cat. no. 20040534) according to manufacturer's guidelines and sequenced with 2x100 bp read length on an Illumina NextSeq1000. Read alignment to reference was done with STAR (Spliced Transcripts Alignment to a Reference).[2] Then, read counts were normalized to log2-counts per million (normalized log2CPM) and integrated in two previously published RNA-Seq data sets[3] : i) 559 bulk RNA-Seq samples of whole bone marrow from B-ALL patients at initial diagnosis, and ii) FACS-sorted B cell populations from four healthy bone marrow donors.[3] All downstream RNA-Seq analyses were done in R Version 4.3.0.[4] Dimensionality reduction (UMAP) was performed with the "umap" page,[5] using the 2802 genes that identify molecular subtype.[3] Graphical visualizations were plotted using ggplot2 package.[6]

To compare gene expression of leukemic blasts with healthy B cell populations, only those patient samples were used where the library preparation and sequencing protocols were identical to rule out batch effects.

#### Cell lines and culture conditions

The B-ALL cell lines 697, SEM, NALM-6, NALM-16 and REH, including its genetically modified versions, were cultured in RPMI 1640 media with 10 % heat-inactivated fetal calf serum (FCS) and 1 % antibiotics, for MHH-CALL-4 and TOM-1 20 % FCS was used. MUTZ-5 cells were cultured with McCoy´s media with 20 % FCS and 1 % antibiotics. Cell lines were incubated in a humidified 5 % $CO_2$ atmosphere at 37 °C. All cells were regularly surveilled for *Mycoplasma* infection using the Venor™GeM Advance detection kit followed by conventional PCR. For in vivo assays, REH cells were subjected to lentiviral transduction with luciferase-GFP ($REH^{LUC}$). Cell lines were authenticated by short tandem repeat (STR) profiling prior to the start of the experiments.

#### Flow cytometry

For analyses of in vitro-cultured cell lines, mouse monoclonal antibodies (mAb) against CD19 (clone 4G7), CD20 (clone 2H7), CD38 (clone HB7), CD47 (clone CC2C6) and against SIRPα (clone 15-414) were purchased from Biolegend. The mAb against total CD47 (clone B6H12) was acquired from Thermo Fisher Scientific. Primary antibodies were incubated for 1 h at 4°C in the dark, and binding was detected with FITC-conjugated goat anti-mouse IgG $F(ab')_2$ fragments (Jackson ImmunoResearch Laboratories). Binding of human antibodies was evaluated using FITC-labelled goat anti-human kappa light chain $F(ab')_2$ fragments (Southern Biotech). Binding of CD47 fusion proteins (50 µg/ml) to M0 macrophages was analyzed by indirect immunofluorescence using FITC-conjugated mouse anti-human IgG $F(ab')_2$ fragments. Fluorophore-conjugated secondary antibodies were incubated for 30 min at 4°C in the dark.

For staining of mouse blood, 50 µl whole blood was stained for 15 minutes at room temperature. Expression of CD89 was determined using anti CD89 (clone A59, Biolegend). The following antibodies were used to determine different effector cells: monocytes were determined using CD45 (clone 30-F11, Biolegend), Ly-6C (clone HK1.4, Biolegend) and CD11b (clone M1/70, Biolegend), granulocytes were identified using CD45, CD11b in combination with Ly-6G (clone 1A8, Biolegend). To determine if the injected anti-CD38 IgA was bound to human CD89 expressed by granulocytes, we stained with a polyclonal $F(ab)_2$ goat anti human IgA (Southern Biotech). To determine if the SIRPα-Fc fusion protein was bound on CD47 expressed by granulocytes, we stained with a polyclonal $F(ab)_2$ goat anti human IgG (Jackson ImmunoResearch). Erythrocytes were lysed for 5-10 minutes at room temerature using freshly made FACSlysings solution (Beckton Dickinson). After a washing step with phosphate buffer saline (PBS), cell pellets were taken up in PBS containing 5 µm latex beads (Molecular Probes, 5000x diluted). The relative amount of effector cells was related to a constant amount of beads (5000 beads)

**Antibody production, purification and quality control**

Purity and integrity of the antibodies were assessed by size exclusion chromatography-high performance liquid chromatography (SEC-HPLC) and SDS-PAGE under reducing conditions using self-prepared 12 % acrylamide gels (running time of 90 min at 120 V, stained with ROTI®Blue staining solution (Carl Roth).

For the therapeutic in vivo study, anti-CD38 IgA2 was produced by WuXi Biologics and validated for size and monomeric state (Suppl. Figure 7A). Cross-validation experiments demonstrated equal binding and efficacy in PMN-mediated ACC matching with the in-house produced anti-CD38 IgA2 (Suppl. Figure 7B, C).

Endotoxin levels of the anti-CD38 IgA2 were < 0.0189 EU/mg. Peptide mapping and glycosylation profiling by liquid chromatography (LC)/mass spectrometry (MS) following MS/MS was supported by GenCC and conducted at Baiya Phytopharm. The mass spectrometry proteomics data have been deposited to the ProteomeXchange Consortium via the PRIDE[7] partner repository with the dataset identifier PXD066660. The raw data will be made accessible once the manuscript has been accepted. Peptide mapping revealed the expected sequences with a coverage of 15.4 % and 100 %, respectively. Glycoprofiling determined two glycosylation sites at N253 and at N349. The first glycosylation site was mainly glycosylated with G0 (44.65%), G1 (32.93%) and G2 (16.29%) glycans, while the second glycosylation site was non-glycosylated (Suppl. Figure 7D).

**Isolation of human effector cells**

PMN and PBMC were isolated from citrate-anticoagulated blood by density gradient centrifugation, utilizing either Polymorphprep® (Progen) or Ficoll Paque Plus (Cytiva). For the generation of non-polarized (M0) macrophages, PBMC were incubated with monocyte attachment medium (PromoCell) for 30 min at 37 °C, followed by three washes with phosphate buffered saline (PBS), and resuspended in X-VIVO™ 15 media (Lonza). After 24 h, 50 ng/ml macrophage colony-stimulating factor (M-CSF) (PeproTech) was added and replenished every 72 h at least twice before cells were used. Fc receptor expression on PMN[8] and M0-like macrophages and the immunophenotype of these macrophages have been described before.[9]

**Antibody-dependent effector cell-mediated assays**

Antibody-dependent cellular phagocytosis (ADCP) with M0 macrophages was measured by live-cell imaging using the IncuCyte® System (Sartorius). B-ALL cells were washed and adjusted to a concentration of $1 \times 10^6$ cells/ml in PBS. Cells were stained with 0.5 µg/ml of the pH-sensitive red fluorescent dye pHrodo (Thermo Fisher Scientific) and incubated in the dark for 1 h at room temperature. Detached macrophages were resuspended in X-VIVO™ 15 media and 20,000 cells per well seeded into a 96-well flat bottom plate (Sarstedt). Antibodies and tumor cells were added at an effector-to-target cell (E:T) ratio of 1:1. Phagocytosis was measured at 37 °C every 20 minutes for a total of 6 h and is defined as red object counts per image (ROI). Analyses were performed using the Incucyte® software (v2019B), with Top-Hat segmentation, using 5 red calibrated units (RCU) and a maximum radius of 20 µm as threshold.

Antibody-dependent PMN-mediated cytotoxicity (ADCC) was measured by chromium-51 [$^{51}$Cr] (Revvity) release assays. Target cells were incubated with $^{51}$Cr for 2 h at 37 °C and washed with cell medium three times. PMN were added at an effector to target cell (E:T) ratio of 80:1 to perform ADCC in the presence of the antibodies of interest and granulocyte-macrophage colony-stimulating factor (GM-CSF, 50 U/ml, CellGenix) for 3 h at 37 °C. $^{51}$Cr release was quantified as counts per minute (cpm) in a MicroBetaTrilux 1450 liquid scintillation and luminescence counter (PerkinElmer). While basal release was measured in the absence of antibodies, maximal $^{51}$Cr release was achieved by addition of 2 % v/v Triton-X 100. Specific tumor cell lysis in percent can be calculated using the following equation:

$$specific\ lysis\ [\%] = \frac{(experimental\ cpm - basal\ cpm)}{(maximal\ cpm - basal\ cpm)}\ x\ 100$$

**In vivo studies**

Mice were transferred from Janvier to the University of Utrecht's Central Laboratory Animal Research Facility, where they were acclimatized for at least 1 week prior to the start of the experiment. In this facility, mice were accommodated in sterile, individually ventilated cages in a regulated environment with a 12:12 h light-dark cycle, and were provided with access to food and water ad libitum.

*Engraftment study:*

10 female CD89 tg SCID and 10 female CD89 tg NXG mice were each injected with 5 x $10^6$ REH$^{LUC}$ cells in the tail vein in 100 µL PBS. Tumor outgrowth was followed by serial bioluminescence imaging (BLI) performing front and back scans 1-2 times per week. (Optical imager, Milabs). BLI images were processed using postprocessing software (Milabs).

*Therapeutic study:*

For the therapeutic study, female NXG mice were used and divided into 6 groups, using CD89 tg or wt mice as indicated for the specific groups (Figure 4A). 4 x $10^6$ REH$^{LUC}$ cells were injected on day 0. All mice received s.c. peg-G-CSF (20 µg/mouse) on days -2 and +5. Mice were randomly assigned to treatment groups using RandoMice (v.1.17) according to genotype. Treatment with anti-CD38 IgA2 (5 mg/kg, i.p.) was given 3 times a week starting at day 0 and continued until day +23. A soluble SIRPα-Fc fusion protein (30 mg/kg, i.p.) was given twice to subgroups of mice. Outgrowth of REH$^{LUC}$ cells was monitored by bioluminescence imaging (BLI) up to 63 days. Front and back scans were taken, and BLI images were processed using postprocessing software (Milabs). Both treatments and analyses were performed in a double-blind manner. Once a week, blood was collected from the submandibular vein in lithium heparin coated tubes (Sarstedt) and used for flow cytometry.

**Generation and characterization of soluble wild-type CD47-Fc and its Q1A variant**

CD47-Fc fusion proteins with a native CD47 ectodomain (CD47-wt; *National Library of Medicine*, NM_001777[55]) and with a mutated CD47 ectodomain (CD47-Q1A variant) were modeled (Figure 5D) using MODELLER and Chimera X software.[10,11] Then, they were generated *de novo* using a pcDNA3.1(+) vector (sequences available upon request) and transient expression in CHO-S cells. Both proteins contain a functionally silent IgG1σ Fc part.[12] After gel electrophoresis, the relevant protein bands (approx. 42 kDa) were analyzed by mass spectrometry (ESI Orbitrap, Q Exactive HF, Thermo Fisher Scientific) after deglycosylation by PNGase and trypsin digestion to identify N-terminal pGlu in CD47-wt or alanine in CD47-Q1A (Figure 5E). The mass spectrometry proteomics data have been deposited to the ProteomeXchange Consortium via the PRIDE[7] partner repository with the dataset identifier PXD066444. The raw data will be made accessible upon the acceptance of the manuscript. Binding of the CD47 fusion proteins to human SIRPα or CD47 antibodies was tested in enzyme-linked immunosorbent assays (ELISA, Figure 5F, Suppl. Figure 6B). Plates were coated with 0,5 µg/ml of the recombinant ectodomain of human SIRPα (SinoBiological) or the CD47 antibodies B6H12 (Thermo Fisher) and CC2C6 (BioLegend). Binding of fusion proteins was detected with goat HRP-conjugated anti-human IgG (Fc-specific) antibodies (ImmunoReagents, 1:5000). Absorption was measured at 450 nm (reference wavelength 540 nm) with Sunrise Basic Microplate Reader (Tecan).

### Supplemental Figures

#### Suppl. Figure 1

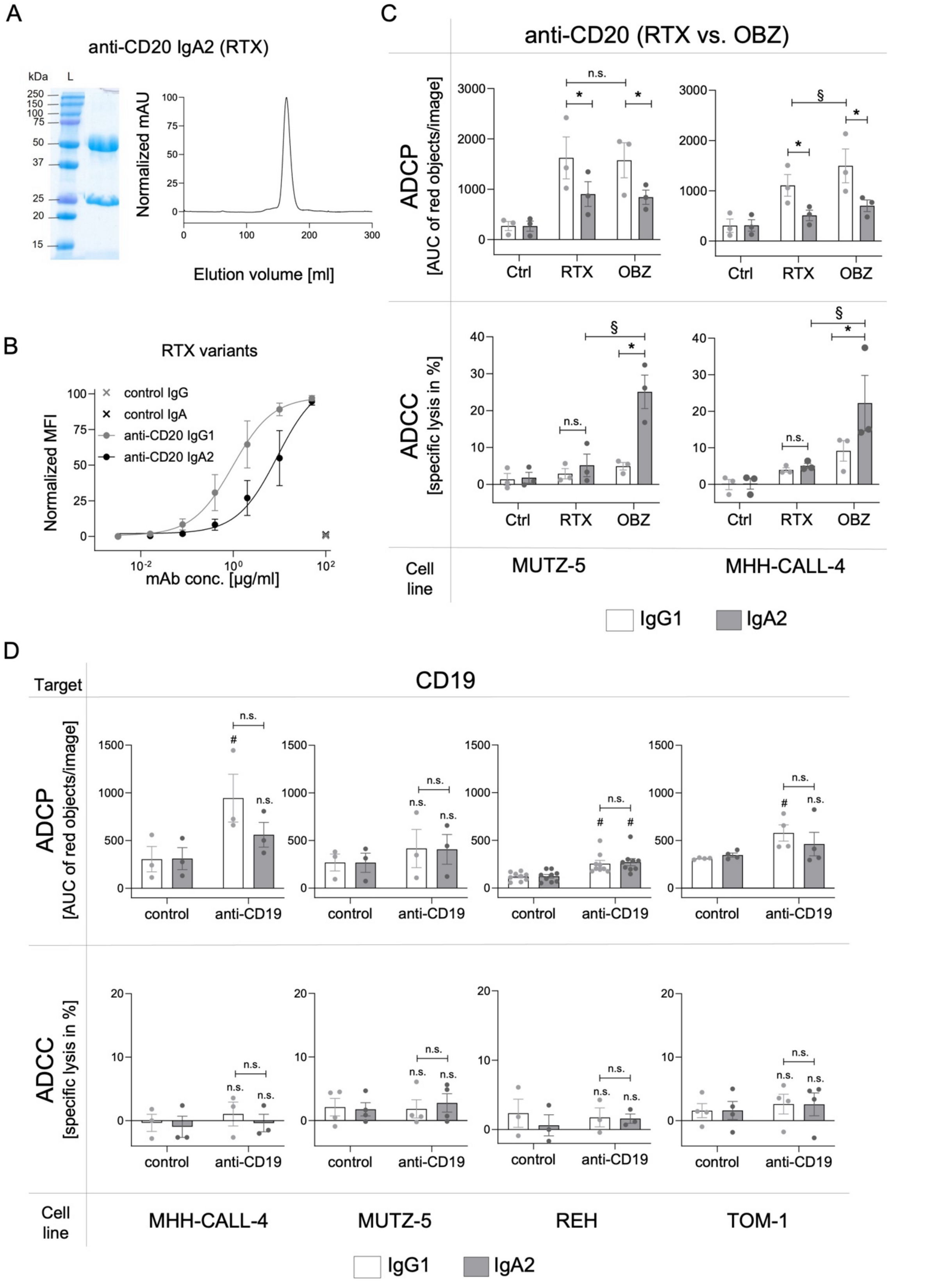


**Suppl. Figure 1. Myeloid cell-mediated tumor cell killing with CD19, CD20, and CD38 antibodies of IgG1 and IgA2 isotypes. (A)** SDS-PAGE under reducing conditions and size exclusion chromatography of rituximab (RTX) IgA2 antibody under native buffer conditions. A representative chromatography image shows the isolated peak fractions normalized to maximum absorption. **(B)** Binding capacities of the RTX IgA2 and RTX IgG1 on MUTZ-5 cells at increasing concentrations up to 50 µg/ml. Antibodies were detected by FITC labelled goat anti-human kappa light chain F(ab')2 fragments. Shown are the normalized MFI values +/- SEM of three independent experiments. **(C)** Comparison of obinutuzumab (OBZ) and RTX IgG1 and IgA2 variants (10 µg/ml) in ADCP and ADCC against B-ALL MUTZ-5 and MHH-CALL-4 cell lines. Data were analyzed by two-way ANOVA, and significant differences ($p \leq 0.05$) between IgG1 and IgA2 (*) and between RTX and OBZ (§) are indicated, n.s., not significant. **(D)** ADCP (upper panel) and ADCC (lower panel) mediated cytotoxicity against four B-ALL cell lines (MHH-CALL-4, MUTZ-5, REH, TOM-1) in the presence of anti-CD19 antibodies (10 µg/ml). Irrelevant IgG1 and IgA2 antibodies were used as controls. Mean values +/- SEM of at least three independent experiments with different donors are indicated as bars, dots represent the values of individual experiments performed in triplicates. Data were analyzed by two-way ANOVA, and significant differences ($p \leq 0.05$) between IgG1 and IgA2 (*) or specific vs control antibodies (#) are indicated. N.s., not significant.

### Suppl. Figure 2

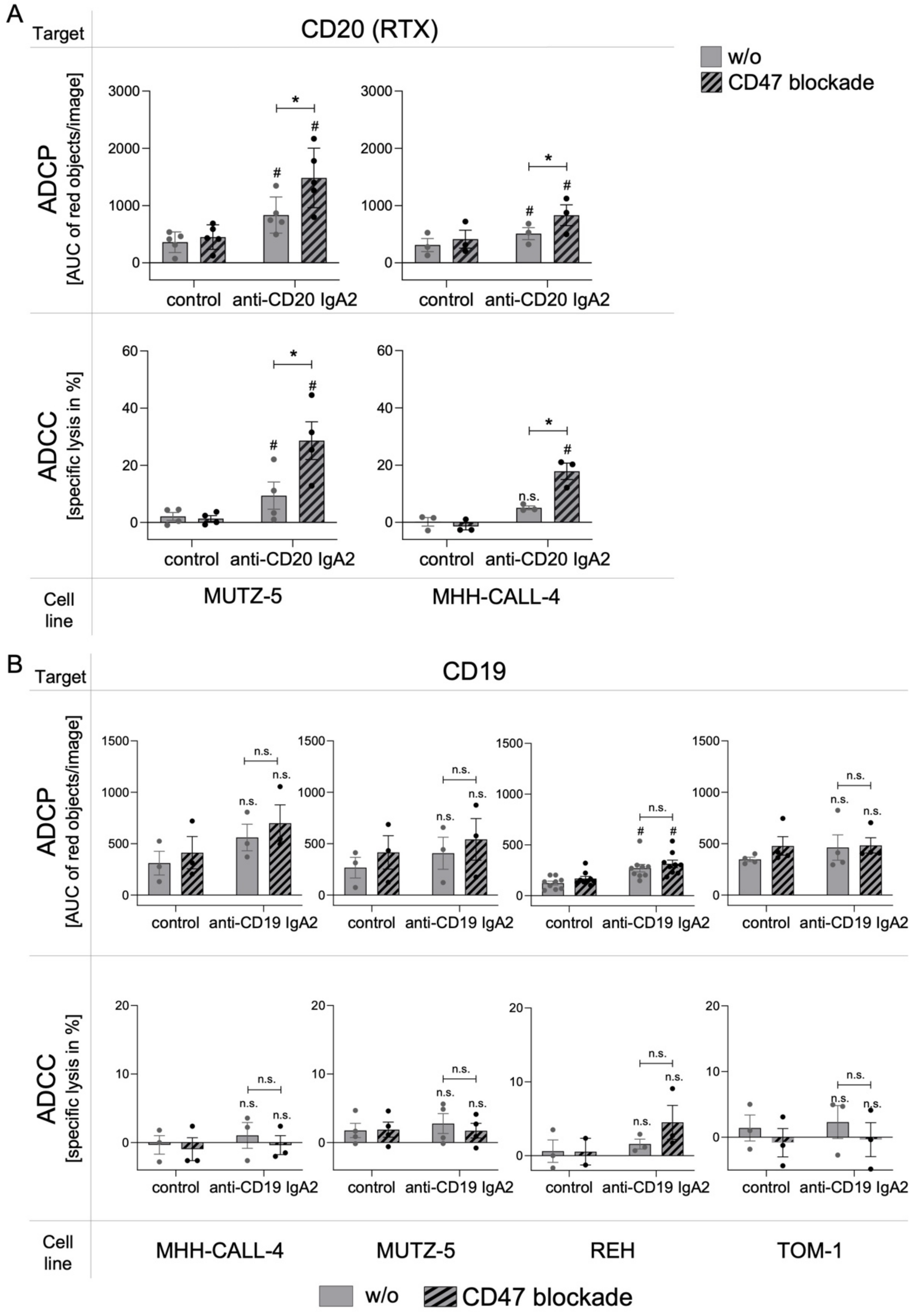

**Suppl. Figure 2. Impact of CD47 blockade on antibody-mediated ADCP and ADCC by IgA2 antibodies against CD20 and CD19.** **(A)** RTX IgA2 (10 µg/ml) alone and in combination with anti-CD47 IgG2s (20 µg/ml) against MUTZ-5 and MHH-CALL-4 in ADCP (upper panels) and ADCC (lower panels). **(B)** Anti-CD19 IgA2 (10 µg/ml) was combined with anti-CD47 IgG2σ (20 µg/ml) in ADCP and ADCC experiments. The assays were performed and are depicted in the same manner as in Suppl. Figure 2A. Results are shown as mean values +/- SEM of at least three independent experiments. Dots represent individual experiments. All data were analyzed by two-way ANOVA, and significant differences ($p \leq 0.05$) between IgA2 alone or in combination with CD47 blockade (*) or specific vs control antibodies (#) are indicated. N.s., not significant.

### Suppl. Figure 3

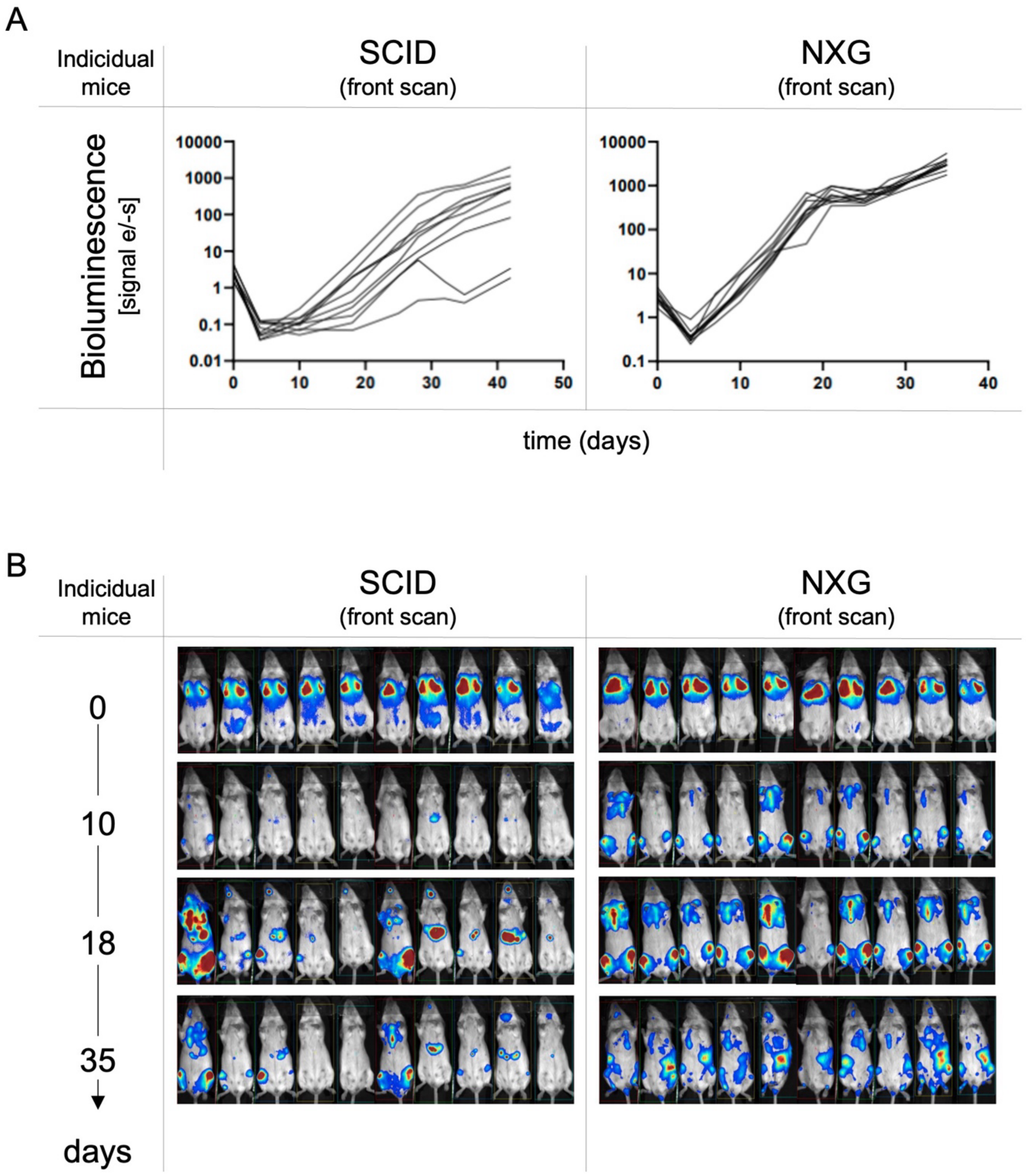


**Suppl. Figure 3. Outgrowth of REH$^{LUC}$ in FcαRI transgenic SCID and NXG mice.** **(A)** Outgrowth of REH$^{LUC}$ cells in SCID and NXG mice over time. Mice were injected with $5x10^6$ REH$^{LUC}$ cells and subjected to bioluminescence (BLI) imaging to follow cancer cell engraftment. BLI data of front scans of individual mice grouped in SCID or NXG are shown. **(B)** Outgrowth of REH$^{LUC}$ cells in SCID and NXG mice over time as presented by front scan BLI images at the indicated days.

### Suppl. Figure 4

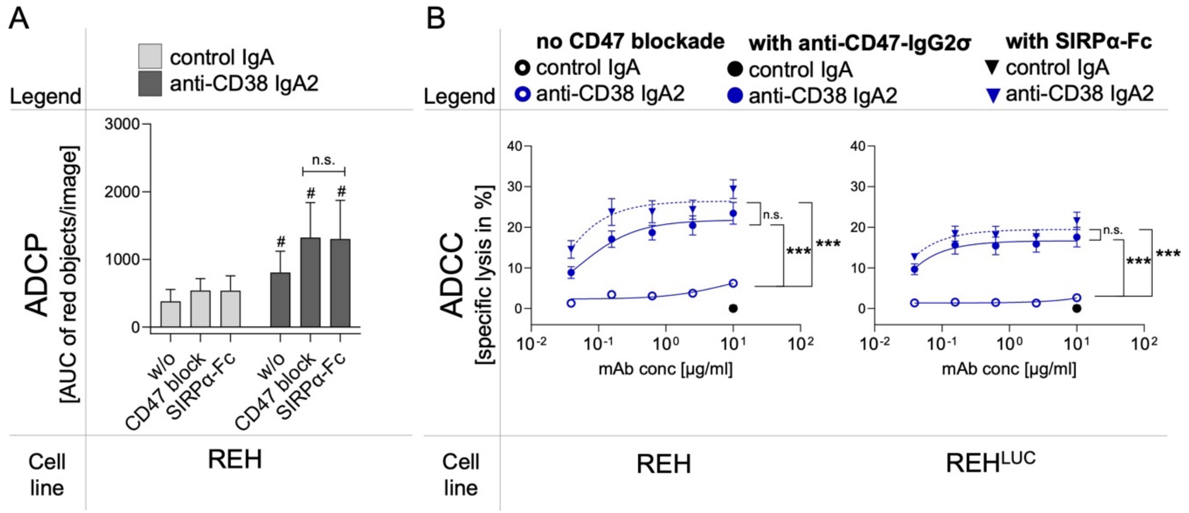


**Suppl. Figure 4. SIRPα-Fc fusion protein used for the in vivo experiment shows similar characteristics and effector functions as the CD47 blocking antibody used for in vitro experiments. (A+B)** The combination of anti-CD38 IgA2 (10 µg/ml) with two different CD47 blocking reagents, CD47 IgGσ antibody or soluble SIRPα-Fc fusion protein (both at 20 µg/ml), was compared in ADCP **(A)** and ADCC **(B)** experiments using REH cells as targets. Shown are mean values +/- SEM of four independent experiments. Data in were analyzed by two-way ANOVA, and significant differences ($p \leq 0.05$) between the anti-CD38 IgA2 alone or in combination with CD47 blocking reagents (*), and specific vs control antibodies (#) are indicated. Differences between the two CD47 blocking reagents were not significant (n.s.).

### Suppl. Figure 5

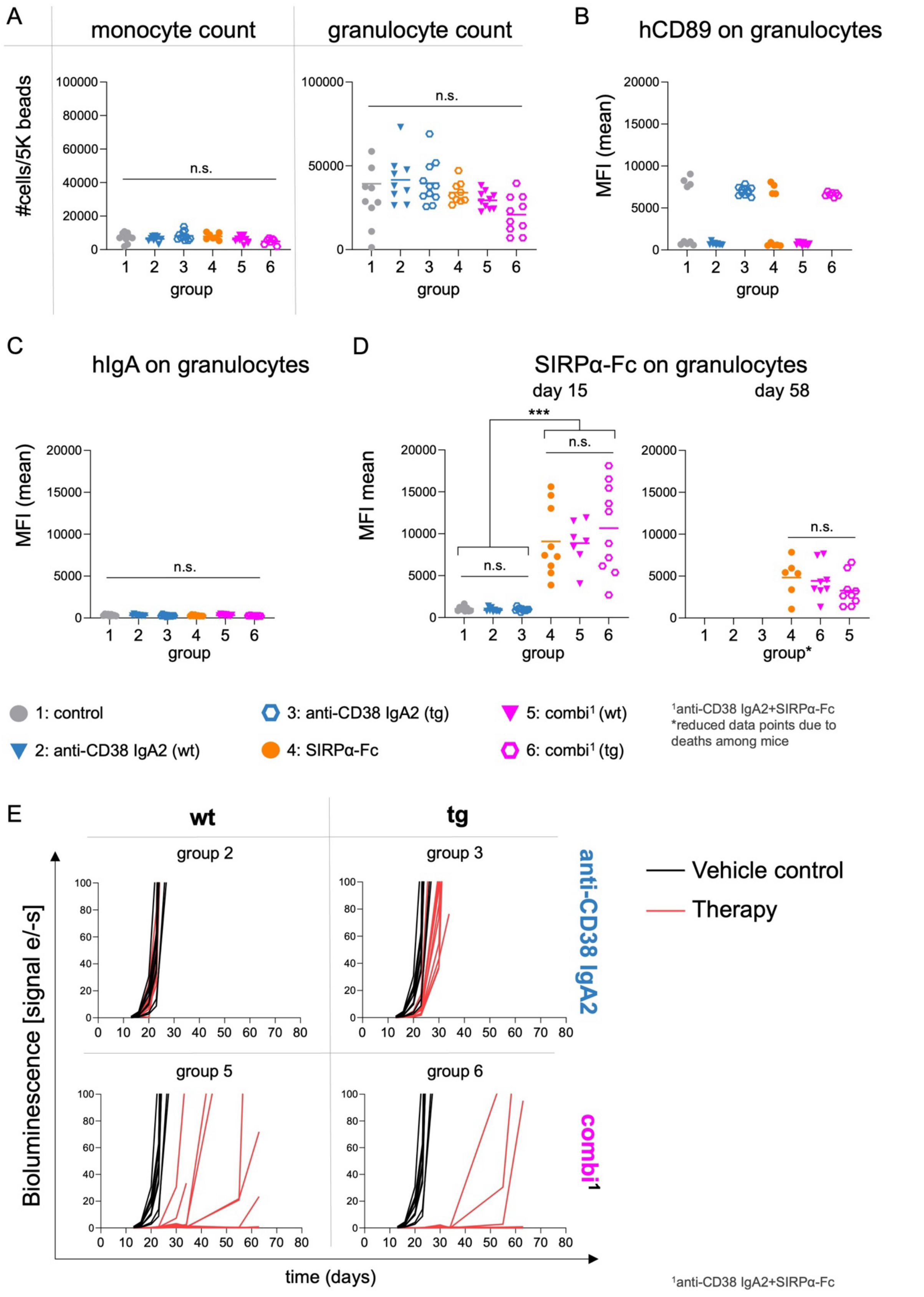


**Suppl. Figure 5. Outgrowth of REH$^{LUC}$ in FcαRI transgenic and wt NXG mice under different therapeutic conditions and flow cytometric analyses of effector cells. (A)** Monocyte (mCD45+/ Ly-6C+/CD11b+) and granulocyte (mCD45+/ Ly-6G+/CD11b+) counts in blood were determined by flow cytometry on day 15 in mice injected with REH$^{LUC}$ cells across the different treatment groups. **(B)** Staining for human CD89, **(C)** human IgA (to detect cytophilic binding of anti-CD38 IgA2), or **(D)** human

IgG (to detect SIRPα-Fc binding) on granulocytes was analyzed by flow cytometry using peripheral blood at the indicated days from different treatment groups. **(E)** Outgrowth of REH$^{LUC}$ cells in individual CD89 transgenic (tg) or wt NXG mice depending on CD89 presence as shown by BLI data from group 4 (anti-CD38 IgA2, wt mice), group 3 (anti-CD38 IgA2, tg mice), group 6 (anti-CD38 IgA2 + SIRPα-Fc [combi], wt mice) and group 5 (anti-CD38 IgA2 + SIRPα-Fc [combi], tg mice).

### Suppl. Figure 6

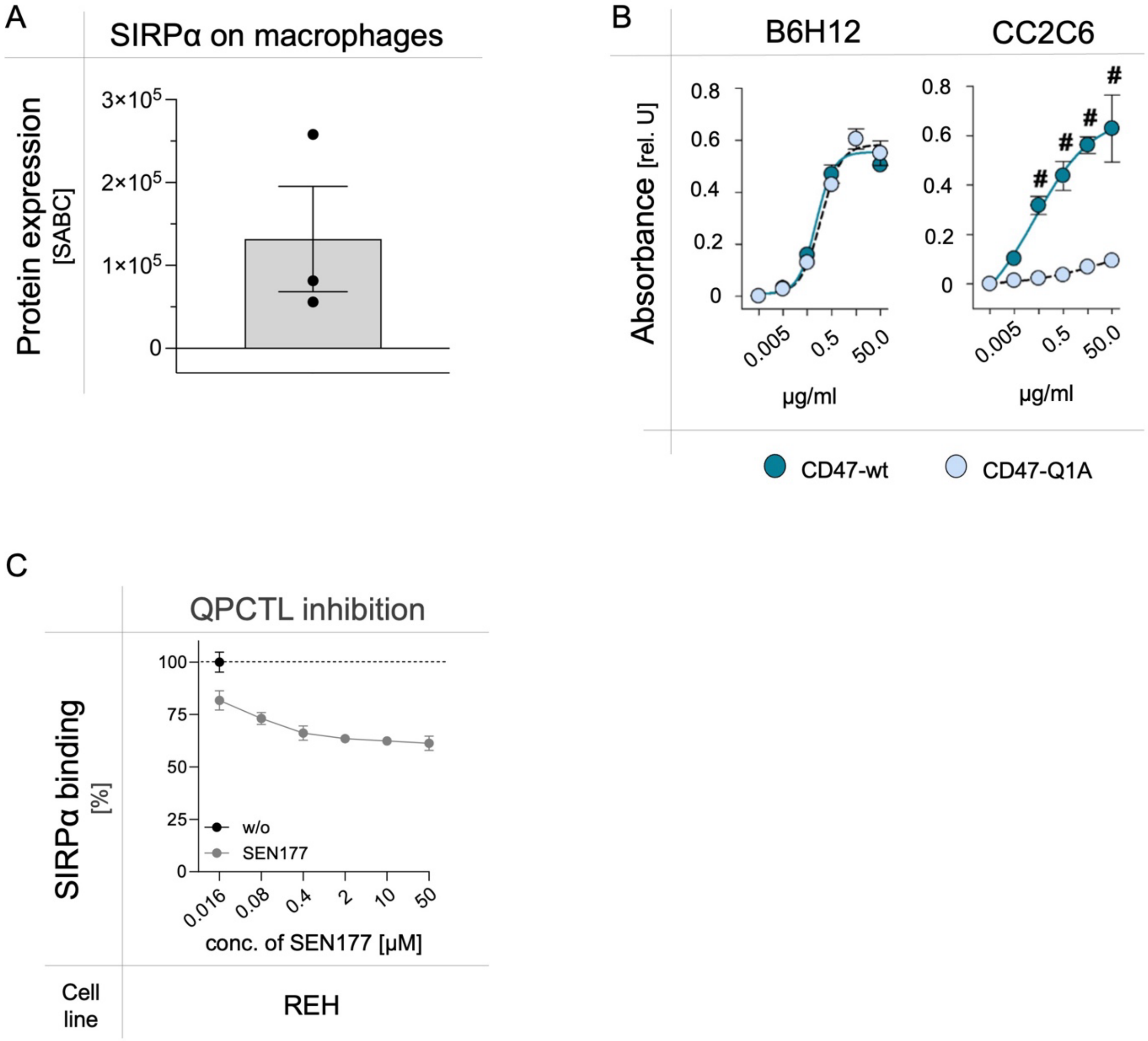


**Suppl. Figure 6. N-terminal pyroglutamate formation of CD47 mediated by QPCTL is important for binding to SIRPα. (A)** SIRPα expression on macrophages was evaluated by indirect flow cytometry using QIFIKIT and a mouse monoclonal SIRPα antibody (clone 15-414, 10 µg/ml), detected by anti-mouse IgG FITC-conjugated F(ab)$_2$ fragments. Macrophages were differentiated from monocytes and results are depicted as mean values ± SEM specific antibody binding capacity (SABC) from three experiments with different donors. **(B)** Differential binding of CD47-wt and CD47-Q1A to recombinant CD47-targeting antibody clones B6H12 (pGlu-independent) and CC2C6 (pGlu-dependent). B6H12 and CC2C6 (each at 0.5 µg/ml) were coated to ELISA plates, CD47 fusion proteins were added at indicated concentrations, and binding detected by mouse HRP-conjugated anti-human IgG (Fc-specific). Mean absorbance values ± SEM of three independent experiments are shown. # indicates statistical differences between CD47-wt and CD47-Q1A at the indicated concentrations ($p < 0.05$ by two-way ANOVA with Bonferroni correction). **(C)** Binding of soluble SIRPα-Fc to REH cells pretreated with increasing concentrations of the QPTCL inhibitor SEN177 for 72 h, DMSO was used as control. Soluble SIRPα-Fc was detected with FITC-conjugated anti-human IgG F(ab')$_2$ fragments.

### Suppl. Figure 7

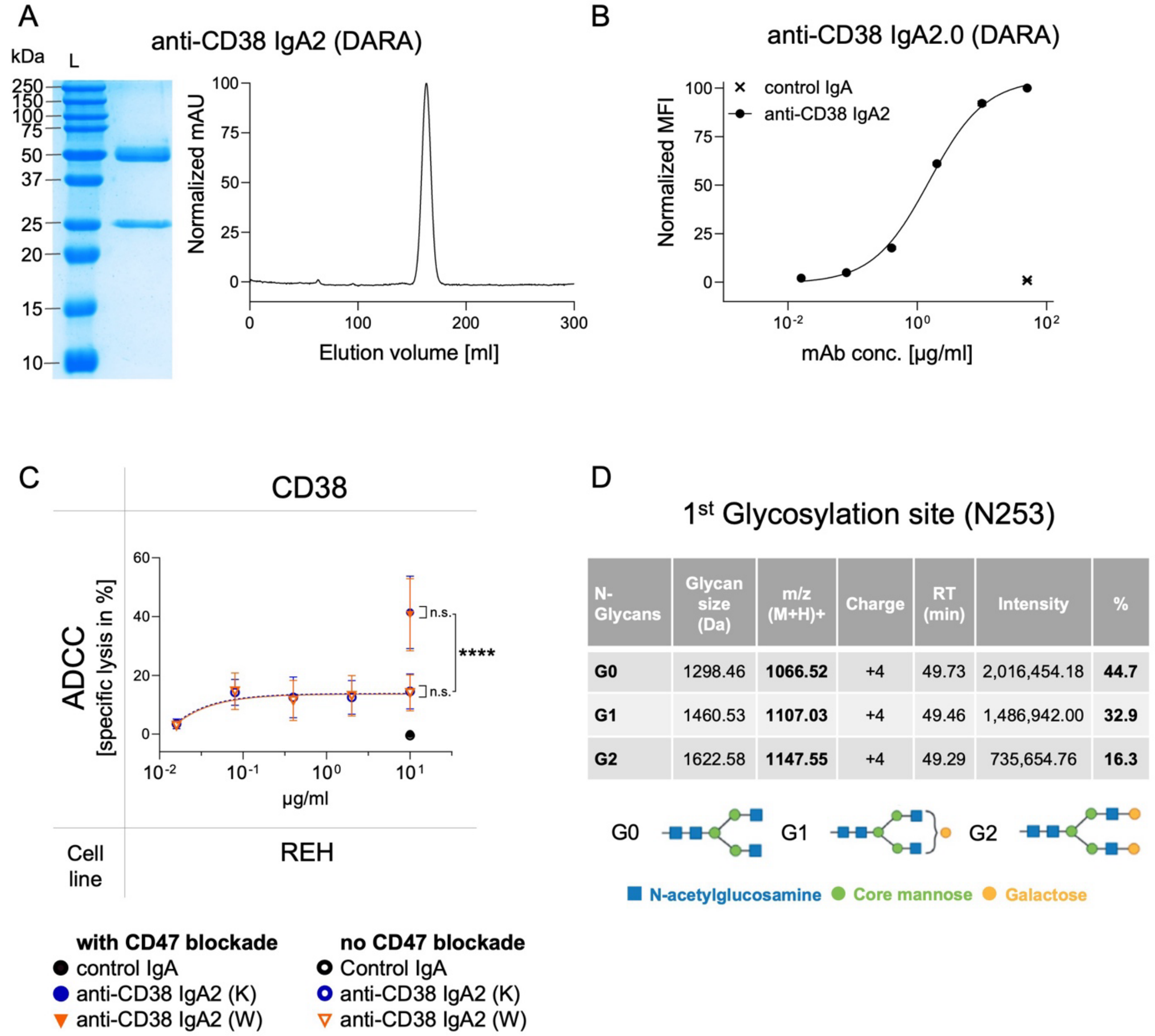


| N-Glycans | Glycan size (Da) | m/z (M+H)+ | Charge | RT (min) | Intensity | % |
|---|---|---|---|---|---|---|
| **G0** | 1298.46 | **1066.52** | +4 | 49.73 | 2,016,454.18 | **44.7** |
| **G1** | 1460.53 | **1107.03** | +4 | 49.46 | 1,486,942.00 | **32.9** |
| **G2** | 1622.58 | **1147.55** | +4 | 49.29 | 735,654.76 | **16.3** |

**Suppl. Figure 7. Anti-CD38 IgA2 produced by WuXi Biologics and used for the in vivo experiment shows similar characteristics and effector functions as the anti-CD38 IgA2 produced in-house and used for in vitro experiments.**
**(A)** Anti-CD38 IgA2, produced by WuXi Biologics, was analyzed by SDS-PAGE under reducing conditions (L, protein ladder) and by size exclusion chromatography under native buffer conditions. **(B)** Binding capacity of the anti-CD38 IgA2 produced by WuXi Biologics on CD38-positive REH cells at increasing concentrations up to 50 µg/ml. The antibody was detected by FITC labelled goat anti-human kappa light chain F(ab')$_2$ fragments. Shown are normalized MFI values +/- SEM of three independent experiments. **(C)** Anti-CD38 IgA2 produced in-house (K) or by WuXi Biologics (W) were tested in ADCC assays at increasing concentrations and combined with anti-CD47 IgG2σ (20 µg/ml) at the highest anti-CD38 IgA2 concentration. Symbols are often overlaying since both antibodies show virtually identical lysis rates. All results represent three independent experiments with effector cells from different donors, each performed in triplicates. **(D)** Glycoprofiles were analyzed by LS/MS following MS/MS. Presented are the three most common N-glycans at the first glycosylation site (N253), while the second site (N449, not shown) was not glycosylated. (RT, retention time).

### References

1. Krohn S, Boje AS, Gehlert CL, Lutz S, Darzentas N, Knecht H, Herrmann D, Brüggemann M, Scheidig AJ, Weisel K, Gramatzki M, Peipp M, Klausz K. Identification of new antibodies targeting malignant plasma cells for immunotherapy by next-generation sequencing-assisted phage display. *Front Immunol*. 2022;13. doi:10.3389/fimmu.2022.908093

2. Dobin A, Davis CA, Schlesinger F, Drenkow J, Zaleski C, Jha S, Batut P, Chaisson M, Gingeras TR. STAR: ultrafast universal RNA-seq aligner. *Bioinformatics*. 2013;29(1):15-21. doi:10.1093/bioinformatics/bts635

3. Beder T, Hansen BT, Hartmann AM, Zimmermann J, Amelunxen E, Wolgast N, Walter W, Zaliova M, Antić Ž, Chouvarine P, Bartsch L, Barz MJ, Bultmann M, Horns J, Bendig S, Kässens J, Kaleta C, Cario G, Schrappe M, Neumann M, Gökbuget N, Bergmann AK, Trka J, Haferlach C, Brüggemann M, Baldus CD, Bastian L. The gene expression classifier ALLCatchR identifies B-cell precursor ALL subtypes and underlying developmental trajectories across age. *Hemasphere*. 2023;7(9):e939. doi:10.1097/HS9.0000000000000939

4. R Core Team. R: A language and environment for statistical computing. Published online 2023. https://www.R-project.org/

5. Konopka T. UMAP: Uniform manifold approximation and projection. Published online February 1, 2023. doi:10.32614/CRAN.package.umap

6. Wickham H. *Ggplot2: Elegant Graphics for Data Analysis*. Springer; 2016. doi:10.1007/978-3-319-24277-4

7. Perez-Riverol Y, Bandla C, Kundu DJ, Kamatchinathan S, Bai J, Hewapathirana S, John NS, Prakash A, Walzer M, Wang S, Vizcaíno JA. The PRIDE database at 20 years: 2025 update. *Nucleic Acids Res*. 2025;53(D1):D543-D553. doi:10.1093/nar/gkae1011

8. Lustig M, Chan C, Jansen JHM, Bräutigam M, Kölling MA, Gehlert CL, Baumann N, Mester S, Foss S, Andersen JT, Bastian L, Sondermann P, Peipp M, Burger R, Leusen JHW, Valerius T. Disruption of the sialic acid/Siglec-9 axis improves antibody-mediated neutrophil cytotoxicity towards tumor cells. *Front Immunol*. 2023;14:1178817. doi:10.3389/fimmu.2023.1178817

9. Lustig M, Hahn C, Leangen Herigstad M, Andersen JT, Leusen JHW, Burger R, Valerius T. Sialylation inhibition improves macrophage mediated tumor cell phagocytosis of breast cancer cells triggered by therapeutic antibodies of different isotypes. *Front Oncol*. 2024;14. doi:10.3389/fonc.2024.1488668

10. Šali A, Blundell TL. Comparative protein modelling by satisfaction of spatial restraints. *J Mol Biol*. 1993;234(3):779-815. doi:10.1006/jmbi.1993.1626

11. Meng EC, Goddard TD, Pettersen EF, Couch GS, Pearson ZJ, Morris JH, Ferrin TE. UCSF ChimeraX: Tools for structure building and analysis. *Protein Sci*. 2023;32(11). doi:10.1002/pro.4792

12. Tam SH, McCarthy SG, Armstrong AA, Somani S, Wu SJ, Liu X, Gervais A, Ernst R, Saro D, Decker R, Luo J, Gilliland GL, Chiu ML, Scallon BJ. Functional, Biophysical, and Structural Characterization of Human IgG1 and IgG4 Fc Variants with Ablated Immune Functionality. *Antibodies (Basel)*. 2017;6(3):12. doi:10.3390/antib6030012
